# Cargo-free scaffold implant recruits metastatic cancer cells *via* lung-mimicking myeloid cell S100A8/A9 axis

**DOI:** 10.1101/789974

**Authors:** Jing Wang, Matthew S. Hall, Grace G. Bushnell, Sophia M. Orbach, Ravi M. Raghani, Yining Zhang, Joseph T. Decker, Aaron H. Morris, Pridvi Kandagatla, Jacqueline S. Jeruss, Lonnie D. Shea

## Abstract

Pre-metastatic niches in distant tissue facilitate metastasis from the primary tumor. Cargo-free porous polymer scaffolds implanted in tumor-bearing mice act as synthetic metastatic niches recruiting metastasizing cancer cells. Herein, we investigated the mechanisms by which these implants attract cancer cells from circulation. Scaffolds attract cancer cells in part *via* S100A8/A9 secreted by Gr1^+^ myeloid cells in a mechanism that mimics lung metastasis. Further, cancer cells attracted to the scaffold have a lung-tropic gene expression signature regardless of their tissue of origin. The scaffold implant reduces metastasis to the lung suggesting otherwise lung-tropic cancer cells are diverted to the scaffold. The suppression of metastatic spread by the scaffold suggests this mechanism may be exploited for novel therapies, and may broadly influence the design of scaffold-based drug delivery system for anti-cancer therapy.

## Introduction

The development of the pre-metastatic niche in native tissues facilitates metastasis by attracting cancer cells from circulation and providing a supportive microenvironment^1-3^. Cargo-free porous polymer scaffolds implanted in hosts bearing breast cancers develop a local microenvironment that functions as a synthetic pre-metastatic niche^4-9^. Synthetic pre-metastatic niches can capture metastasizing cancer cells before they are detectable in native organs^5, 6^, show potential as a therapeutic by reducing metastatic tumor burden in native organs^6^, and can be engineered and humanized to uniquely study human pathways in native pre-metastatic niche progression^6-9^. Previous work has shown that the detection of cancer cells at the synthetic pre-metastatic niche occurs soon after the arrival of myeloid cells^5, 6^, and identifying the mechanism governing recruitment would improve their application as models^8, 9^ for study of native pre-metastatic niches and as local delivery systems for immunotherapy^10, 11^.

Here, we dissected the contribution of different cell subsets at the synthetic pre-metastatic niche—a cargo-free microporous poly(ε-caprolactone) (PCL) scaffold implant—to cancer cell recruitment. Further, we investigated the common myeloid cell secreted S100A8/A9 recruitment axis between the synthetic pre-metastatic niche and pre-metastatic lung, which may divert metastasis of lung-tropic cancer cells to the implant. These studies will determine the extent to which cargo-free synthetic pre-metastatic niches mimic the organotropic metastasis observed in native tissues where cancer cells originating in a given organ will preferentially metastasize to metastatic niches in a particular organ ^12, 13^.

## Results and discussion

### Diseased conditioning enhances breast cancer cell recruitment from the blood to scaffold implants

We initially investigated the role of scaffold size and number on cancer cell recruitment (Fig. 1A and Supplementary Figs 1 and 2). Cargo-free porous polymer scaffolds recruit metastasizing cancer cells from the primary tumor in orthotopic 4T1 murine breast cancer models that develop spontaneous metastasis^5, 6^. Interestingly, a large scaffold implant recruited substantially more metastasizing cancer cells than a small scaffold (Fig. 1A and Supplementary Fig. 1), yet the ratio of cancer cells to all cells at the implant remained the same regardless of scaffold size (Supplementary Fig. 2). We also found that the number of cancer cells captured per scaffold did not vary with the number of implants per mouse (Fig. 1A), suggesting that the local microenvironment of each scaffold in tumor-bearing mice was responsible for cancer cell recruitment.

**Fig. 1.**
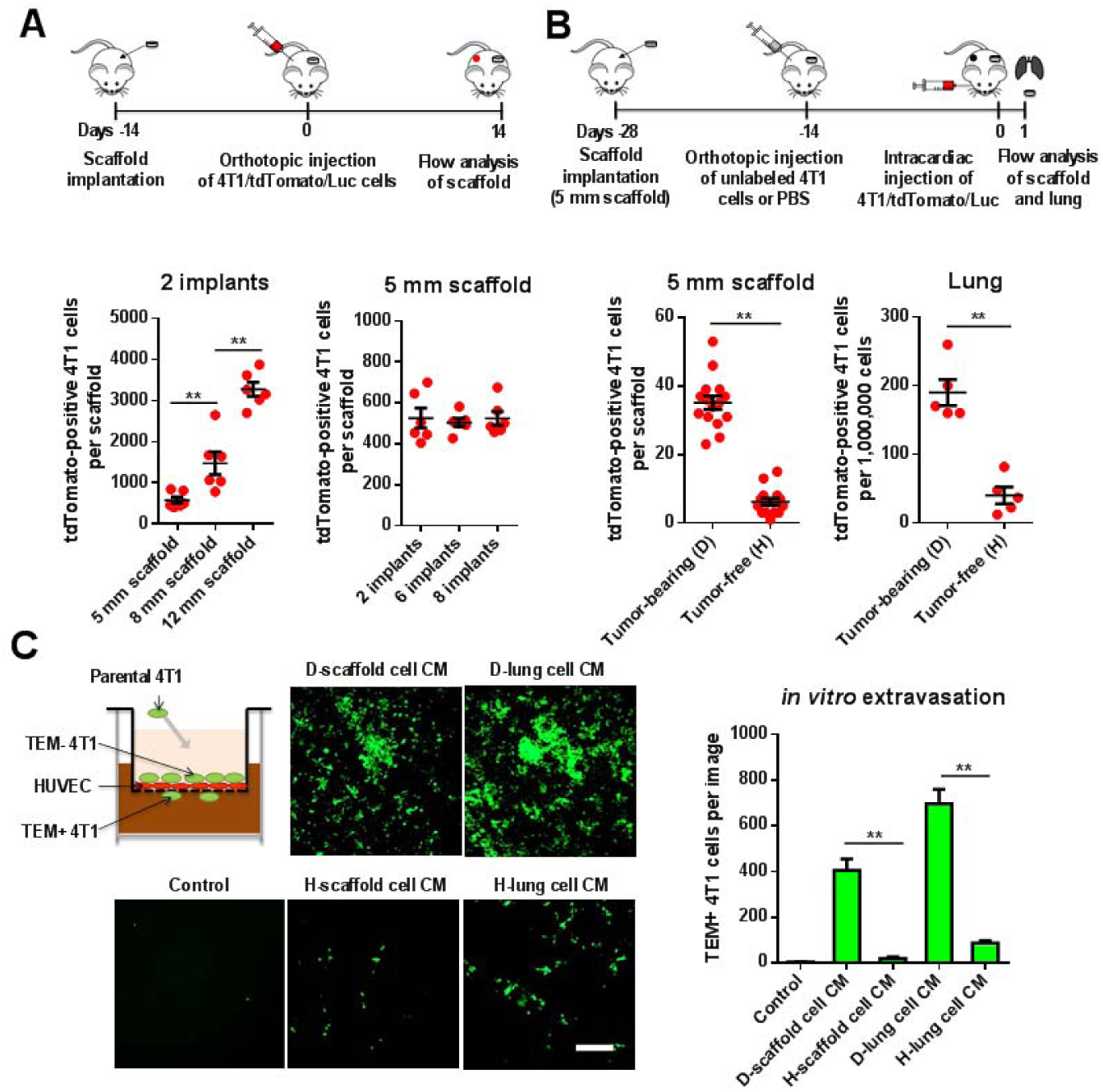
Metastatic breast cancer cells colonize cargo-free porous PCL scaffold implants and the pre-metastatic lung in tumor-bearing mice. (A) Orthotopic spontaneous metastasis model. Scaffold implants recruit tdTomato-labeled 4T1 murine breast cancer cells that spontaneously metastasize from the primary tumor. The number of cancer cells recruited per scaffold is dependent on scaffold size but independent of implant number per mouse. Scaffold implants were 5 mm, 8 mm or 12 mm in diameter. (B) Intracardiac injection extravasation model. tdTomato-labeled 4T1 cancer cells introduced into circulation by intracardiac injection were more numerous in the lungs and scaffold of mice bearing unlabeled 4T1 tumor (“diseased” D) than tumor-free mice (“healthy” H) after 1 day. (C) *In vitro* extravasation of 4T1 cancer cells towards conditioned media (CM) prepared by scaffolds and lungs retrieved from tumor-bearing mice (“diseased” D) or tumor-free mice (“healthy” H). The cancer cells that migrate across the HUVEC monolayer and recover from the lower chamber represent the metastatic cells and are defined as transmigrated cells (TEM+ cells) while the less aggressive cancer cells that stay in the upper chamber are TEM-cells. Conditioned media prepared with cells from diseased scaffolds and lung stimulated more transendothelial migration of 4T1 cancer cells than conditioned media produced with cells from healthy mice. Scale bar = 150 µm. \*\**p* < 0.01, from student *t*-test. All experiments were performed three times with similar results.

The role of diseased conditioning of the scaffold on cancer cell recruitment was next analyzed by performing intracardiac injections^14^ of tdTomato-labeled 4T1 cancer cells (4T1/tdTomato/Luc cells) in scaffold-bearing BALB/c mice that either had an unlabeled 4T1 cell primary tumor (diseased) or no tumor (healthy). After 24 hours, we observed that 5-fold more tdTomato-positive 4T1 cells had migrated from the blood into the scaffolds and lungs of the tumor-bearing mice than the tumor-free mice (Fig. 1B and Supplementary Fig. 3). These *in vivo* results suggest that cells in the diseased scaffolds and lungs may secrete factors that actively induce the extravasation of 4T1 cells from the blood. This hypothesis was tested with media conditioned by cells from diseased and healthy scaffolds and lungs, with quantification of 4T1 cell transmigration towards the conditioned media in an *in vitro* extravasation model (Fig. 1C). The diseased conditioned media of both the scaffold and lung stimulated more 4T1 cells transmigration than the conditioned media produced from healthy tissue (Fig. 1C).

### Expanded granulocytic myeloid cells at the diseased scaffold implant secrete S100A8/A9 to recruit breast cancer cells

We next sought to determine which cell populations at the scaffold secreted the soluble factors that increased 4T1 cell transmigration to diseased scaffolds and lungs. Using flow cytometry, we observed a 5 to 6-fold increase in myeloid cells (CD11b^+^Gr1^+^) in the scaffolds and lungs of diseased mice compared to healthy mice (Fig. 2A, B and Supplementary Fig. 4). These myeloid cells were predominately granulocytic (CD11b^+^Gr1^+^Ly6G^+^) rather than monocytic (CD11b^+^Gr1^+^Ly6C^+^). Macrophages (CD11b^+^F4/80^+^) and dendritic cells (CD11c^+^F4/80^-^) were reduced in the diseased scaffolds and lungs while no statistically significant changes were observed in other cell types at the scaffold. We next prepared conditioned media from multiple diseased scaffold cell populations using Magnetically Assisted Cell Sorting (MACS) on various surface markers. We found that cell subpopulations that included granulocytic myeloid cells (CD45^+^ leukocytes, Gr1^+^ myeloid cells, and Ly6G^+^ granulocytic myeloid cells) greatly enhanced transmigration of 4T1 cells in our *in vitro* extravasation model (Fig. 2C and Supplementary Fig. 5).

**Fig. 2.**
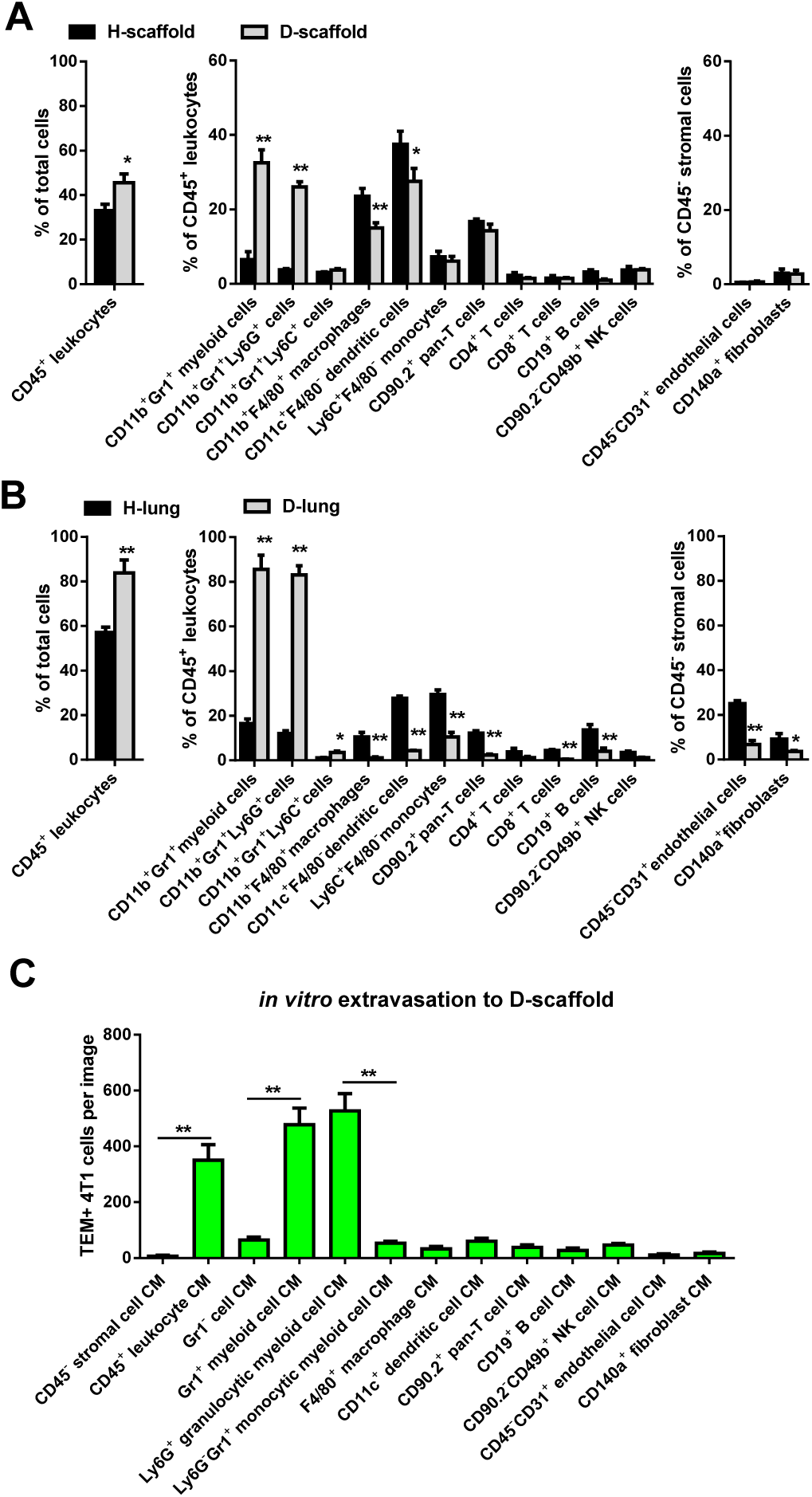
Expanded Gr1^+^ myeloid cells in diseased scaffold implants are responsible for recruiting breast cancer cells. (A, B) Flow cytometry analysis reveals granulocytic myeloid cells (CD11b^+^Gr1^+^Ly6G^+^) are more abundant at the scaffold and lungs of “diseased” day 14 tumor-bearing mice (D) than “healthy” tumor-free mice (H). (C) 4T1 cancer cells exhibit enhanced extravasation *in vitro* towards conditioned media (CM) prepared with cell subpopulations from diseased scaffolds that contain granulocytic myeloid cells (CD45^+^ leukocytes, Gr1^+^ myeloid cells, and Ly6G^+^ granulocytic myeloid cells). The surface markers displayed for each cell subset indicate the antibodies used in magnetic-activated cell sorting. NK: natural killer; TEM: transendothelial migration. \**p* < 0.05, \*\**p* < 0.01, from student *t*-test. All experiments were performed three times with similar results.

We also found that S100A8/A9 mRNA expression and secreted protein in the diseased scaffold and lung were greatly upregulated relative to healthy tissue (Fig. 3A, B). Previous work has shown that myeloid cell secreted S100A8/A9 is a potent chemoattractant for metastasizing cancer cells at the pre-metastatic lung^15, 16^. At the diseased scaffold, S100A8/A9 mRNA and secreted protein was primarily from cell subpopulations containing granulocytic myeloid cells (CD45^+^ leukocytes, Gr1^+^ myeloid cells, and Ly6G^+^ granulocytic myeloid cells) (Fig. 3C, D). Just as in the scaffold, Gr1^+^ myeloid cells in the lung were responsible for most S100A8/A9 mRNA expression and protein secretion (Fig. 3E, F). Neutralizing S100A8/A9 bioactivity in Gr1^+^ myeloid cell conditioned media with an anti-S100A8/A9 antibody eliminated the increase in 4T1 cell transmigration for both lung and diseased scaffold conditions (Fig. 3G and Supplementary Fig. 6). We also demonstrated that the activity of the anti-S100A8/A9 antibody was specific to S100A8/A9 by observing that it reduced 4T1 cell transmigration towards soluble recombinant S100A8/A9 protein (Fig. 3G and Supplementary Fig. 6), but not towards 0.1% fetal bovine serum (FBS) (control, Supplementary Fig. 7).

**Fig. 3.**
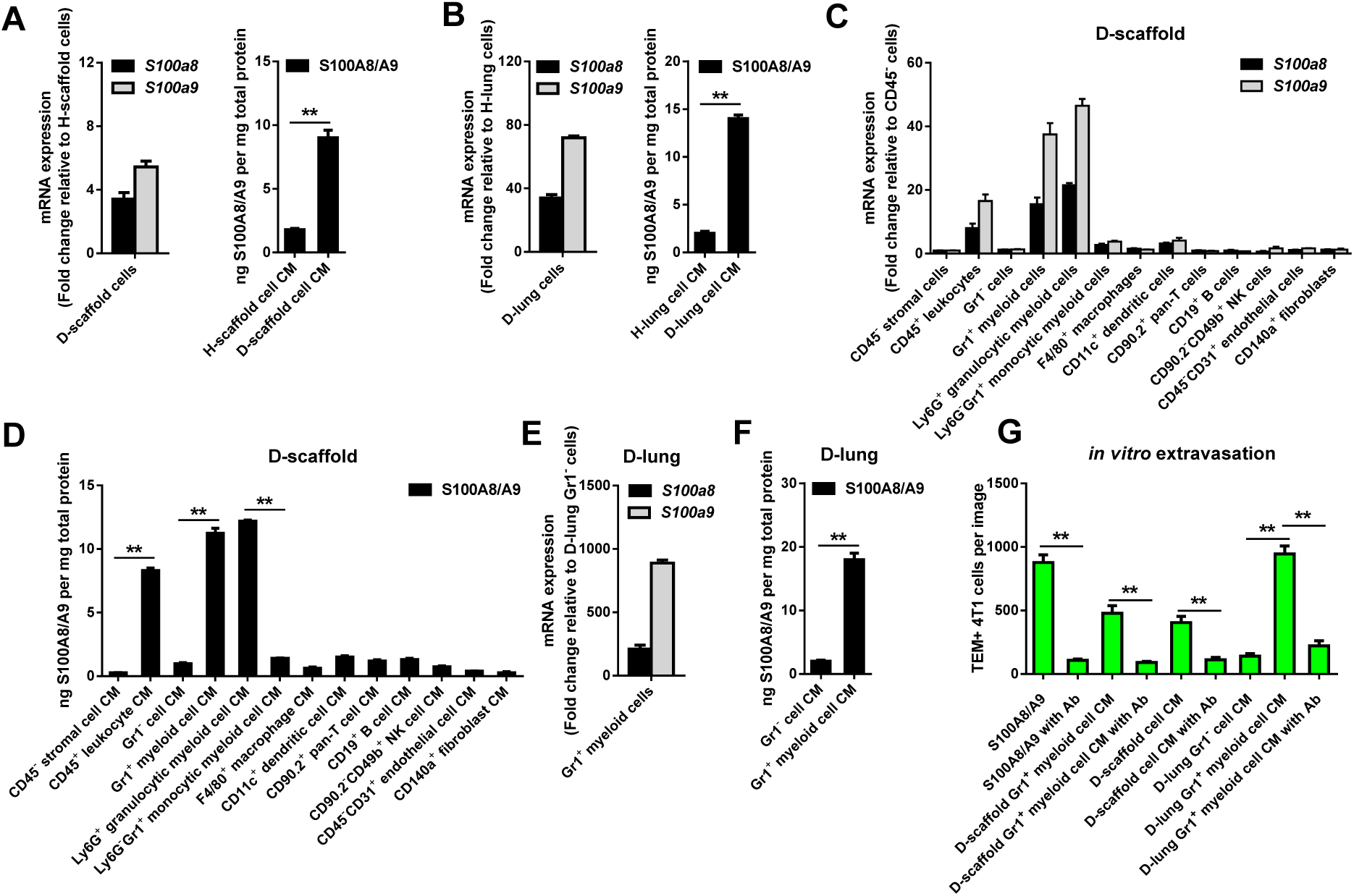
S100A8/A9 secreted by Gr1^+^ myeloid cells activates extravasation of breast cancer cells to diseased scaffolds. (A, B) Gene expression and protein secretion of S100A8/A9 are up-regulated in diseased scaffolds and lungs relative to healthy tissue. (C, D) Gene expression and protein secretion of S100A8/A9 is predominately in cell subpopulations that contain granulocytic myeloid cells (CD45^+^ leukocytes, Gr1^+^ myeloid cells, and Ly6G^+^ granulocytic myeloid cells). (E, F) S100A8/A9 is overexpressed and secreted by Gr1^+^ myeloid cells in diseased lung. (G) Anti-S100A8/A9 antibody inhibits enhanced transmigration of 4T1 cancer cells towards recombinant S100A8/A9 protein (positive control) and conditioned media prepared by Gr1^+^ myeloid cells from diseased scaffold or lung. H: healthy; D: diseased; NK: natural killer; CM: conditioned media; TEM: transendothelial migration; Ab: anti-S100A8/A9 antibody. \*\**p* < 0.01, from student *t*-test. All experiments were performed three times with similar results.

### S100A8/A9 induces a lung-tropic gene signature in breast cancer cells attracted to the scaffold implant

We next compared the gene expression of transmigrated 4T1 cells (TEM+ cells) *in vitro* to those that did not transmigrate (TEM-cells) (Fig. 1C, schematic) using these known gene expression signatures as a metric of induced organotropism. Previous reports have identified differential gene expression signatures of breast cancer cell lines derived from metastases in the lung^17^ and bone^18^ relative to the parent line. We found that diseased lung conditioned media and diseased scaffold conditioned media induced a lung-tropic gene expression signature in transmigrated 4T1 cells, as did soluble recombinant S100A8/A9 protein in isolation (Fig. 4A). By comparison, diseased bone marrow conditioned media and soluble CXCL12, a dominant chemoattractant in bone chemotaxis^19^, induced a bone-tropic gene expression signature in transmigrated 4T1 cells. Two non-organ-specific chemoattractants, CCL2 and 0.1% FBS, induced neither signature in transmigrated 4T1 cells (Fig. 4A). Autocrine signaling through S100A8/A9 is known to amplify myeloid cell recruitment and increase S100A8/A9 at the pre-metastatic lung^15, 16, 20^. Interestingly, we found that initiating 4T1 cancer cells may participate in similar autocrine signaling to recruit following 4T1 cells. 4T1 cell populations that were first to transmigrate towards disease scaffolds, diseased lung, or S100A8/A9 protein overexpressed *Rage, S100a8*, and *S100a9* (Fig. 4A). Subsequent transmigrating cancer cells had lower expression of these genes, which diluted the overall difference between transmigrated and non-transmigrated 4T1 cells (Supplementary Fig. 8).

**Fig. 4.**
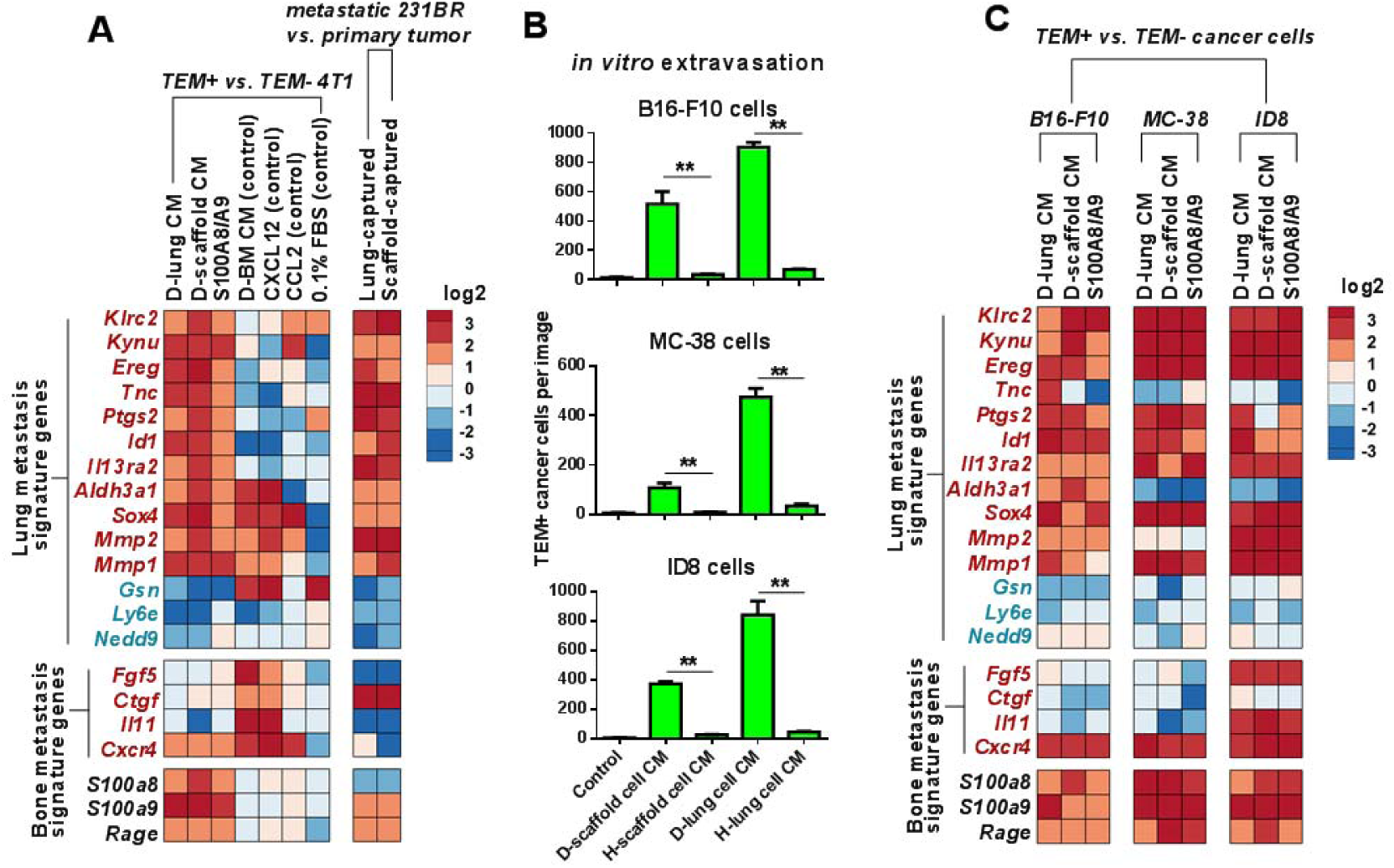
Scaffold-recruited cancer cells have lung-tropic gene expression signature *in vitro* and *in vivo*. (A) Scaffold-homing cancer cells have gene expression more similar to a known lung-tropic^17^, than a known bone-tropic^18^ signature (gene signature names to the left, red text overexpressed in signature, blue text under-expressed in signature). 4T1 cells were allowed to transmigrate towards conditioned media *in vitro* and the relative gene expression of transmigrated (TEM+) to non-transmigrated (TEM-) cancer cells is displayed. Metastasized MDA-MB-231BR (231BR) human breast cancer cells were isolated from the lung and scaffold of tumor-bearing NSG mice. The relative gene expression of lung and bone derived 231BR cell lines compared to a line from the primary tumor is displayed. (B) *In vitro* B16-F10 murine melanoma cells, MC-38 murine colon cancer cells and ID8 murine epithelial ovarian cancer cells all extravasate more extensively towards conditioned media prepared from diseased scaffold or lung relative to healthy tissue. \*\**p* < 0.01, from student *t*-test. (C) Heatmap of log2 fold change of organotropism signature genes in transmigrated (TEM+) B16-F10, MC-38 and ID8 cancer cells relative to their respective non-gransmigrated (TEM-) cell lines show lung-tropic gene expression for cells attracted to the diseased scaffold. H: healthy; D: diseased; CM: conditioned media; TEM: transendothelial migration; BM: bone marrow; FBS: fetal bovine serum. All experiments were performed three times with similar results.

Our attempts to isolate and expand 4T1 cells directly from diseased lung and scaffold tissue for validation of these results *in vivo* were unsuccessful due to technical limitations. However, we were able to isolate cancer cells from diseased lung and scaffold using MDA-MB-231BR human breast cancer cells^21^. Both the diseased lung and scaffold isolated cancer cell lines aligned with the lung-tropic gene expression signature, but not the bone-tropic signature, demonstrating the lung-like recruitment mechanism towards the cargo-free scaffold implants *in vivo* (Fig. 4A). We also observed *Rage* and *S100a9* overexpression in these cell lines relative to the parent line, suggesting that the cancer cell S100A8/A9 autocrine loop might be present at the synthetic pre-metastatic niche *in vivo* (Fig. 4A).

Collectively, these results show that the key chemoattractant in the pre-metastatic niche, whether native or synthetic, can selectively recruit the metastatic cancer cell subpopulation with the specific phenotype associated with organotropic metastasis. This finding suggests that synthetic pre-metastatic niches that model different organotropisms could be engineered by altering their material chemistry, delivering cells, or the sustained release of a soluble chemoattractant^4-9^.

### Cells from diseased scaffolds attract lung-tropic cancer cell subpopulation from different tissues of origin

Lung metastasis is common in cancers across many tissues of origin^22^, and we next investigated whether the lung-tropic recruitment axis to the diseased scaffold was unique to 4T1 breast cancer cells or applied more universally to other cancers. B16-F10 (murine skin melanoma), MC-38 (murine colon), and ID8 (murine ovarian) cancer cell lines all transmigrated more extensively to diseased scaffold and lung conditioned media than healthy (Fig. 4B and Supplementary Fig. 9). As was the case for the 4T1 breast cancer cell line, the differential gene expression signature of transmigrated cells aligned with a lung-tropic rather than bone-tropic signature (Fig. 4C). These results suggest that the diseased scaffold induces a common and universal lung-tropic cellular phenotype in recruited cancer cells similar to that found for cancer cells recruited to the lung^17^.

In aggregate, these results show that cargo-free porous polymer scaffolds act as synthetic metastatic niches that mimic the lung’s S100A8/A9 recruitment axis. As in the lung, myeloid cells at the scaffold implant secrete soluble S100A8/A9 to recruit metastasizing cancer cells, resulting in that scaffold-recruited cancer cells express a lung-tropic gene expression signature regardless of their tissue of origin.

### Scaffold implants suppress growth of recruited cancer cells and reduce lung metastasis

Metastatic cancer cells with a lung-tropic gene expression signature can colonize the pre-metastatic lung and scaffold implants, yet only the lung supported their outgrowth. An *in vitro* study indicated that media condition by total cells or the Gr1^+^ myeloid cell subset harvested from the diseased scaffold inhibited 4T1 cancer cell growth by causing apoptosis of cancer cells (Fig. 5A and Supplementary Fig. 10) in a dose dependent manner, while similar conditioned media from diseased lung enhanced growth (Fig. 5A). Unlike the lung, metastatic cancer cells that colonized the scaffold implant did not exhibit outgrowth into secondary tumors (Fig. 5B, C and Supplementary Figs 11-15).

**Fig. 5.**
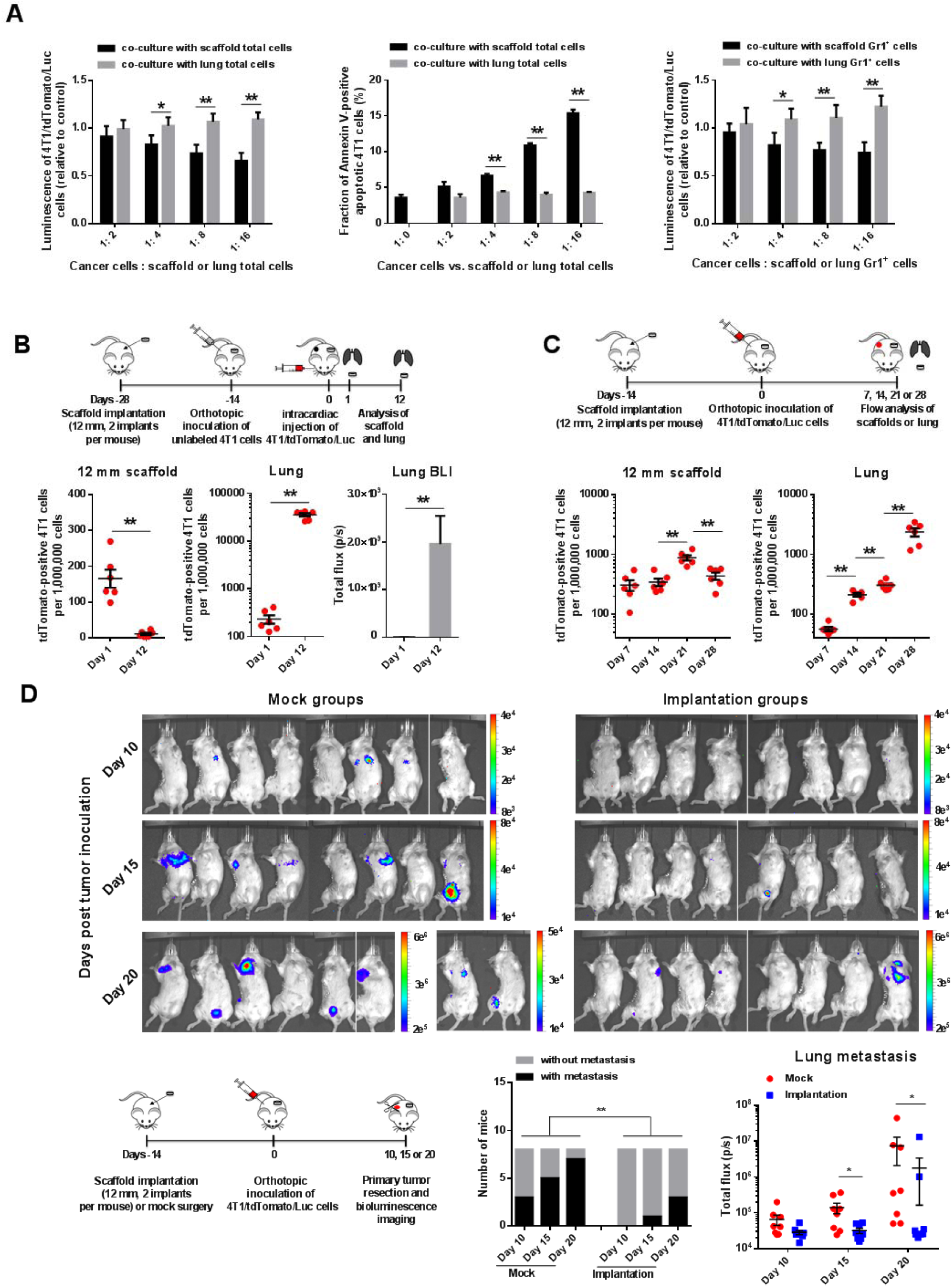
Scaffold implants suppress growth of locally recruited cancer cells and reduce metastasis to the lung. (A) *In vitro*, co-culture of 4T1/tdTomato/Luc cancer cells with cells from the diseased scaffolds (total cells or Gr1^+^ myeloid cell subpopulation) inhibits the proliferation of 4T1 cells and causes cancer cell apoptosis while similar cell populations from diseased lung promote 4T1 cell growth. Cancer cells were co-cultured with lung or scaffold cells for two days in complete cell culture media and cancer cells without co-culture with other cells were used as control. \**p* < 0.05 and \*\**p* < 0.01, from student *t*-test. (B) *In vivo*, in the intracardiac injection extravasation model, 4T1/tdTomato/Luc cancer cells that extravasated to the scaffold from circulation fail to outgrow, decreasing in abundance from days 1 to 12 after intracardiac injection. In contrast, tdTomato-positive cancer cells in the lung proliferate and expand over this time. 4T1/tdTomato/Luc cancer cells were introduced into circulation by intracardiac injection in BALB/c mice bearing unlabeled 4T1 primary tumor, and 1 day or 12 days later, scaffold implants and lungs were retrieved. The abundance of tdTomato-positive cancer cells in these tissues was quantified by flow cytometry. Increased bioluminescence of the Luc-positive 4T1 cells in the lungs of mice was also detected *ex vivo*. BLI: bioluminescence imaging. \*\**p* < 0.01, from student *t*-test. (C) *In vivo*, in the orthotopic spontaneous metastasis model, 4T1/tdTomato/Luc cancer cells that metastasized to the scaffold implants from the primary tumor fail to outgrow, decreasing in abundance from days 21 to 28 while fluorescent cancer cells in the lung massively increase over this time. Scaffold implants and lungs were retrieved from mice bearing 4T1/tdTomato/Luc tumor at the days 7, 14, 21 and 28 after tumor inoculation and the tdTomato-positive cancer cells in these tissues were analyzed by flow cytometry. \*\**p* < 0.01, from student *t*-test. (D) Scaffold implantation suppresses lung metastasis. Mice were implanted with 12 mm PCL scaffolds (2 implants per mouse, implantation groups) or received mock surgery at the same location (mock groups), and two weeks later, six groups (n = 8 mice per group) were inoculated with 2×10^6^ of 4T1/tdTomato/Luc cancer cells. For reference, the average humane endpoint of mice in this model is ∼28 days without treatment. At days 10, 15 or 20 after tumor inoculation, one mock group and one implantation group received primary tumor resection to allow for more sensitive *in vivo* bioluminescence imaging of metastasis, and then the bioluminescence of metastatic cancer cells was immediately imaged. Mice bearing scaffold implants were less likely to have detectable metastasis than mice that only received mock implantation surgery. \*\**p* < 0.01, from Fisher’s exact test. Mice bearing scaffolds also had reduced luminescence localized to the lung in the day 15 and day 20 groups (Supplementary Fig. 16). \**p* < 0.05, from the non-parametric Mann Whitney test.

We first examined whether 4T1 cells that trafficked to the scaffold implant remained viable or died sometime after reaching the implant. We used our intracardiac injection model to introduce 4T1/tdTomato/Luc cells into circulation in mice bearing unlabeled 4T1 primary tumors (Fig. 5B, schematic). In this model, the bolus of tdTomato and Luc labeled 4T1 cancer cells introduced into circulation by intracardiac injection is clear within 1 to 2 days^23^. We therefore quantified the abundance of 4T1/tdTomato/Luc cells in the scaffolds and lungs 1 day and 12 days after intracardiac injection. While we observed far more 4T1/tdTomato/Luc cells in the lungs 12 days after intracardiac injection than 1 day (Fig. 5B and Supplementary Figs 11 and 12), indicating massive proliferation, there were far fewer 4T1/tdTomato/Luc cells at the scaffold on day 12 than on day 1 (Fig. 5B and Supplementary Figs 11 and 13), indicating lack of proliferation and cell death. These results strikingly demonstrate a tumoricidal property of the synthetic metastatic niche implant toward metastatic cancer cells that may contribute to its utility as a potential therapeutic.

In BALB/c mice that were initially orthotopically inoculated with fluorescent 4T1/tdTomato/Luc cancer cells (Fig. 5C, schematic), cancer cells metastasized to the scaffold implant from primary tumor by 7 days after inoculation (Fig. 5C and Supplementary Figs 14 and 15). Accumulation of cancer cells at the scaffold reached a peak at day 21, possibly due to abundant circulating tumor cells in the blood at this time point. By day 28, the number of cancer cells per scaffold (Supplementary Fig. 15a) as well as the ratio of cancer cells to total cells at the scaffold (Fig. 5C and Supplementary Fig. 14) had decreased despite the total cells recovered per scaffold remaining constant across time (Supplementary Fig. 15b). In contrast, the cancer cells that colonized the lung massively expanded from days 21 to 28, developing into macrometastases by day 28 (Fig. 5C), necessitating endpoint for the experiment. These results further support the conclusion that the microenvironment of the scaffolds does not support tumor cell outgrowth.

In aggregate, these results suggest that lung-tropic metastatic cancer cells that would otherwise metastasize to the lung are attracted to the synthetic pre-metastatic niche *via* S100A8/A9 and are prevented from forming secondary tumors in the scaffold implants. Further supporting this mechanism, we observed that scaffold implants suppressed the development of lung metastasis without influencing the growth of primary tumor in the orthotopic 4T1 breast cancer model (Fig. 5D and Supplementary Figs 16 and 17). Additional studies that investigate the mechanisms underlying the differential outgrowth within the lung and scaffold may inspire novel therapeutics for metastasis.

Our findings are broadly useful across the emerging field of scaffold-based local delivery system for anti-cancer therapy^10, 11^, including in the design of vaccine^24^, drug^25^ and cell^26^ carrying therapeutic implants. The accumulation of S100A8/A9 secreting myeloid-derived cells and the arrival of metastatic cancer cells might alter the function of the payloads in these systems. For example, Gr1^+^ myeloid cells may suppress the trafficking and anti-cancer activity of biomaterial-delivered T cells^27^ while cancer cells that colonize the scaffold implant might stimulate the antigen-presentation of dendritic cells together with scaffold-delivered cancer vaccines^28^. Cargo-free polymer scaffolds may also find utility as a direct therapy, capturing and eliminating lung-tropic cancer cells from circulation that would otherwise metastasize to the lung (Fig. 5) and therefore potentially improve the survival after tumor resection for patients with aggressive metastatic disease^6^. Further studies establishing how the base-line response to biomaterial scaffold implants in tumor-bearing hosts differ from those in healthy individuals^29^ would serve as a foundation for engineering better scaffold-based drug delivery systems for anti-cancer therapy^10, 11,30^.

## Acknowledgements

This research was supported by the National Institutes of Health (R01CA214384). Thanks to Dr. Weiping Zou at the University of Michigan for kindly providing B16-F10 murine melanoma cell line, MC-38 murine colon cancer cell line and ID8 murine epithelial ovarian cancer cell line.

## Author contributions

L.D.S., J.S.J., J.W. and M.S.H. designed the experiments. J.W. conducted all the *in vitro* experiments. J.W. and M.S.H. performed all the *in vivo* experiments. G.G.B. isolated and expanded the human MDA-MB-231BR cells from diseased lung and scaffold and provided technical assistance with intracardiac injection. R.M.R. and Y.Z. helped in studying Gr1^+^ myeloid cells. S.M.O. provided technical assistance with flow cytometry. J.T.D., A.H.M. and P.K. provided laboratory assistance and suggestions for the experiments. L.D.S., J.S.J., J.W. and M.S.H. wrote and edited the manuscript.

## Additional information

Supplementary information is available for this paper. Correspondence and requests for materials should be addressed to L.D.S. and J.S.J.

## References

1. Kaplan, R. N., Rafii, S. & Lyden, D. Preparing the “soil”: The premetastatic niche. Cancer Res 66, 11089–11093 (2006).

2. Joyce, J. A. & Pollard, J. W. Microenvironmental regulation of metastasis. Nat Rev Cancer 9, 239–252 (2009).

3. Liu, Y. & Cao, X. Characteristics and significance of the pre-metastatic niche. Cancer Cell 30, 668–681 (2016).

4. Bersani, F. et al. Bioengineered implantable scaffolds as a tool to study stromal-derived factors in metastatic cancer models. Cancer Res 74, 7229–7238 (2014).

5. Azarin, S. M. et al. In vivo capture and label-free detection of early metastatic cells. Nat Commun. 6:8094 (2015).

6. Rao, S. S. et al. Enhanced survival with implantable scaffolds that capture metastatic breast cancer cells *in vivo*. Cancer Res 76, 5209–5218 (2016).

7. Carpenter, R. A., Kwak, J. G., Peyton, S. R. & Lee, J. Implantable pre-metastatic niches for the study of the microenvironmental regulation of disseminated human tumour cells. Nat Biomed Eng 2, 915–929 (2018).

8. Aguado, B. A., Bushnell, G. G., Rao, S. S., Jeruss, J. S. & Shea, L. D. Engineering the pre-metastatic niche. Nat Biomed Eng 1, 0077 (2017).

9. Beri, P. et al. Biomaterials to model and measure epithelial cancers. Nat Rev Mater 3, 418–430 (2018).

10. Cheung, A. S. & Mooney, D. J. Engineered materials for cancer immunotherapy. Nano Today 10, 511–531 (2015).

11. Weiden, J., Tel, J. & Figdor, C. Synthetic immune niches for cancer immunotherapy. Nat Rev Immunol 18, 212–219 (2018).

12. Obenauf, A. C. & Massagué, J. Surviving at a distance: organ specific metastasis. Trends in cancer 1, 76–91 (2015).

13. Peinado, H. et al. Pre-metastatic niches: organ-specific homes for metastases. Nat Rev Cancer 17, 302–317 (2017).

14. Saxena, M. & Christofori, G. Rebuilding cancer metastasis in the mouse. Mol Oncol 7, 283–296 (2013).

15. Hiratsuka, S., Watanabe, A., Aburatani, H. & Maru, Y. Tumour-mediated upregulation of chemoattractants and recruitment of myeloid cells predetermines lung metastasis. Nat Cell Biol 8, 1369–1375 (2006).

16. Hiratsuka, S. et al. The S100A8–serum amyloid A3–TLR 4 paracrine cascade establishes a pre-metastatic phase. Nat Cell Biol 10, 1349–1355 (2008).

17. Minn, A. J. et al. Genes that mediate breast cancer metastasis to lung. Nature 436, 518–524 (2005).

18. Kang, Y. et al. A multigenic program mediating breast cancer metastasis to bone. Cancer Cell 3, 537–549 (2003).

19. Weilbaecher, K. N., Guise, T. A. & McCauley, L. K. Cancer to bone: a fatal attraction. Nat Rev Cancer 11, 411–425 (2011).

20. Srikrishna, G. S100A8 and S100A9: new insights into their roles in malignancy. J Innate Immun 4, 31–40 (2011).

21. Bushnell, G. G. et al. Biomaterial scaffolds recruit an aggressive population of metastatic tumor cells *in vivo*. Cancer Res 79, 2042–2053 (2019).

22. Budczies, J. et al. The landscape of metastatic progression patterns across major human cancers. Oncotarget 6, 570–583 (2014).

23. Basse, P., Hokland, P., Heron, I. & Hokland, M. Fate of tumor cells injected into left ventricle of heart in BALB/c mice: role of natural killer cells. J Natl Cancer Inst 80, 657–665 (1998).

24. Hu, Z. et al. Towards personalized, tumour-specific, therapeutic vaccines for cancer. Nat Rev Immunol 18, 168–182 (2018).

25. Park, C. G. et al. Extended release of perioperative immunotherapy prevents tumor recurrence and eliminates metastases. Sci Transl Med 10, eaar1916 (2018).

26. Stephan, S. B. et al. Biopolymer implants enhance the efficacy of adoptive T-cell therapy. Nat Biotech 33, 97–101 (2015).

27. Ostrand-Rosenberg, S. et al. Cross-talk between myeloid-derived suppressor cells (MDSC), macrophages, and dendritic cells enhances tumor-induced immune suppression. Semin Cancer Biol 22, 275–281 (2012).

28. Tacken, P. J. et al. Dendritic-cell immunotherapy: from *ex vivo* loading to *in vivo* targeting. Nat Rev Immunol 7, 790–802 (2007).

29. Anderson, J. M., Rodriguez, A. & Chang, D. T. Foreign body reaction to biomaterials. Semin Immunol 20, 86–100 (2008).

30. Wang, H. & Mooney, D. J. Biomaterial-assisted targeted modulation of immune cells in cancer treatment. Nat Mater 17, 761–772 (2018).

