## Supplementary Information for "Cargo-free scaffold implant recruits metastatic cancer cells *via* lung-mimicking myeloid cell S100A8/A9 axis"

**This PDF file includes:**

Materials and Methods
Supplementary Figs 1-18
References for SI reference citations

**Materials and Methods**

Materials……………………………………………………………………….…3
Cell culture…………………………………………………………….…...…...3
Scaffold fabrication and implantation……….……………………….……………..3
Orthotopic injection spontaneous metastasis model………..…………………….…..…..3
Intracardiac injection metastasis model……………………………………….….……3
Cell subset isolation…………………………………………………….................4
Condition media preparation…………………………….…………………….….4
*in vitro* extravasation of cancer cells……………………………………………...4
Immunostaining and flow cytometry…………………………..……………4
Quantitative RT-PCR (qRT-PCR)…………………………………………….…..5
ELISA……………………………………………………………………….….5
Isolation of transmigrated cancer cells from Boyden chamber…………….………….6
Isolation of metastatic cancer cells from orthotopic cancer model……………………....6
*in vitro* viability of 4T1/tdTomato/Luc cancer cells………………………….……...6
*in vivo* bioluminescence imaging………………………………….………...6
Statistical analysis……………………………………………………………...7
**Supplementary Fig. 1.** Representative density plots for tdTomato-positive 4T1 cancer cells that metastasized from the primary tumor to the scaffold implants with different sizes in orthotopic injection spontaneous metastasis model…………………………………..…..8
**Supplementary Fig. 2.** Relative abundance of tdTomato-positive 4T1 cancer cells to total cells in scaffold implants with different sizes (a) and total cells per scaffold (b)..………………..…9

**Supplementary Fig. 3.** Representative density plots for tdTomato-positive 4T1 cancer cells that extravasated to 5 mm scaffold implants and lungs in intracardiac injection metastasis model……………………………………………………………………………..…...10

**Supplementary Fig. 4.** Representative density plots for CD11b$^+$Gr1$^+$ myeloid cells in scaffold implants and lungs……………………………………………………………………….11

**Supplementary Fig. 5.** Microscopy images of 4T1 cancer cells that transmigrated towards conditioned media prepared from different diseased scaffold cell subsets………………12

**Supplementary Fig. 6.** Microscopy images of 4T1 cancer cells that transmigrated towards 20 ng/ml S100A8/A9 (positive control) or conditioned media in the absence or presence of anti-S100A8/A9 antibody…………………………………………………………..13

**Supplementary Fig. 7.** Anti-S100A8/A9 antibody does not inhibit FBS-induced 4T1 cell transmigration ……………………………………………………………………...14

**Supplementary Fig. 8.** The initiating 4T1 cancer cells that transmigrated towards D-lung conditioned media overexpressed *Rage* and *S100a9* genes relative to later transmigrating cells…………………………………………………………………………………15

**Supplementary Fig. 9.** Microscopy images of transmigrated B16-F10 murine melanoma cells, MC38 murine colon cancer cells and ID8 murine epithelial ovarian cancer cells………16

**Supplementary Fig. 10.** Representative density plots for apoptotic cell subpopulation in 4T1 cancer cells co-cultured with scaffold or lung total cells……………………………..17

**Supplementary Fig. 11**. Representative density plots for tdTomato-positive 4T1 cancer cells that extravasated to 12 mm scaffold implants and lungs 1 and 12 days after intracardiac injection of 4T1/tdTomato/Luc cells in mice bearing unlabeled 4T1 tumor……………18

**Supplementary Fig. 12.** Bioluminescence imaging of lungs 1 and 12 days after intracardiac injection of 4T1/tdTomato/Luc cells in mice bearing unlabeled 4T1 tumor…………….19

**Supplementary Fig. 13**. tdTomato-positive 4T1 cells per scaffold (a) and total cells per scaffold (b) 1 and 12 days after intracardiac injection of 4T1/tdTomato/Luc cells in mice bearing unlabeled 4T1 tumor………………………………………………………………….20

**Supplementary Fig. 14**. Representative density plots for tdTomato-positive 4T1 cancer cells that metastasized to scaffold implants and lungs at different time points post orthotopic inoculation of 4T1/tdTomato/Luc cells…………………………………….....……..21

**Supplementary Fig. 15.** tdTomato-positive 4T1 cells per scaffold (a) and total cells per scaffold (b) at days 7, 14, 21 and 20 after orthotopic inoculation of 4T1/tdTomato/Luc cells.…..22

**Supplementary Fig. 16.** The ROI area used to calculate the total flux of lung metastasis in Fig 5D……………………………………………………………………………………...23

**Supplementary Fig. 17.** Averaged primary tumor size at days 10, 15 and 20 after tumor inoculation………………………………………………………………………..…….24

**Supplementary Fig. 18.** A high ratio of TEM+ cells to TEM- cells obscured their gene expression difference ..………………………………..………………………….…..25

**References for SI reference citations**……………………………………………...…………26

### Materials and Methods.

**Materials.** The 4T1 murine breast cancer cell line and human umbilical vein endothelial cell (HUVEC) were purchased from ATCC. The 4T1/tdTomato/Luc murine breast cancer cell line was obtained from Perkin Elmer. The B16-F10 murine skin melanoma cell line, MC-38 murine colon adenocarcinoma cell line, and the ID8 murine ovarian cancer cell line were kindly provided by Dr. Weiping Zhou (University of Michigan). The MDA-MB-231BR (231BR) human breast cancer cell line was obtained from the Northwestern University Developmental Therapeutics Core. Recombinant mouse S100A8/A9 heterodimer (carrier-free) (#765504) and recombinant mouse CCL2 (MCP-1) (carrier-free) (#578402) were purchased from Biolegend. Mouse CXCL12/SDF-1α (#SRP4388) was from Sigma. Anti-S100A8/A9 complex antibody (#ab22506) was from Abcam. All kits and microbeads used for MACS cell separation were purchased from Miltenyi Biotec. Primers were from RealTimePrimers. Mouse S100A8/A9 heterodimer DuoSet ELISA kits were from R&D Systems. CellTracker Green CMFDA dye were purchased was Invitrogen. D-luciferin was from PerkinElmer. Alexa Fluor 488-labeled Annexin V was from ThermoFisher Scientific. All the other chemicals were from Sigma.

**Cell culture.** 4T1 and 4T1/tdTomato/Luc murine breast cancer cells were cultured in RPMI-1640 medium (Gibco) supplemented with 10% heat inactivated fetal bovine serum (FBS). MDA-MB-231BR (231BR) human breast cancer cells, B16-F10 murine skin melanoma cells and MC38 murine colon adenocarcinoma cells were cultured in Dulbecco's modified MEM (DMEM) (Gibco) supplemented with 10% FBS. ID8 murine ovarian cancer cells were cultured in high glucose DMEM (Gibco) supplemented with 4% FBS, 5 µg/ml insulin, 5 µg/ml transferrin and 5 ng/ml sodium selenite (Sigma). HUVEC was cultured in EGM-2 medium (Lonza) with 10% FBS. All the cells were grown at 37 °C with 5% of $CO_2$ and full humidify.

**Scaffold fabrication and implantation.** Microporous poly(caprolactone) (PCL) scaffolds were fabricated by salt leaching technique. Briefly, PCL microspheres were prepared as described previously.[1] Then 0.2 g of PCL microspheres and 6 g sodium chloride salt particles (250-425 µm) were mixed well and pressed at 1500 psi for 45 seconds in a 40 mm diameter die. The polymer/salt discs were heated with a 60 °C hotplate for 5 min on each side, followed by leaching out the salt with distilled water. The polymer discs were dried on kimwipes and then frozen at -80 ºC for more than 1 h. The frozen 40 mm diameter polymer discs were placed on dry ice and cut by the Miltex disposable biopsy punch (5 mm, 8 mm or 12 mm in diameter) to prepare PCL scaffolds (~2 mm in height). Before implantation, the microporous scaffolds were disinfected by soaking in 70% ethanol for 1 minute, rinsed with sterile water, and air dried on a sterile gauze pad. Microporous scaffolds were implanted in the subcutaneous space of the back of either female BALB/c or female NOD-scid IL2Rgammanull (NSG) mice (8-10 weeks old) as described previously.[1-3]

**Orthotopic injection spontaneous metastasis model.** Scaffolds were subcutaneously implanted in female BALB/c mice (3-6 mice per group). Two weeks later, $2\times10^6$ fluorescent 4T1/tdTomato/Luc cancer cells in 50 µl of PBS was inoculated into the fourth right mammary fat pad by injection. At the desired day post tumor inoculation, scaffolds and lungs were retrieved and prepared to single cell suspension as described before,[1,2] and the tdTomato-positive tumor cell number in the scaffold and lungs was quantified by flow cytometer (Bio-Rad ZE5 Cell Analyzer or MoFlo Astrios #2 Cell Sorter).

**Intracardiac injection metastasis model**. Scaffolds were implanted in female BALB/c mice (8-10 weeks). After two weeks, either $2\times10^6$ non-fluorescent 4T1 cancer cells in 50 µl of PBS (tumor-bearing, 3-6 mice per group) or 50 µl PBS without cells (tumor-free, control) was inoculated into the fourth right mammary fat pad by injection. At day 14 post tumor inoculation, a total of $1\times10^6$ fluorescent 4T1/tdTomato/Luc cells in 100 µl of PBS were injected into the left ventricle of both tumor-bearing mice (diseased, D) and tumor-free mice (healthy, H). And one day later, mice were euthanized and the lungs and scaffolds were retrieved, processed to single cell suspension as described previously,[1,2] and measured by flow cytometer to identify the number of tdTomato-positive cancer cells in lungs and scaffolds.

**Cell subset isolation.** Subsets of lung or scaffold cells were isolated by positive selection with magnetic-activated cell sorting (MACS). First, a single cell suspension of cells from scaffolds and lungs was prepared as described previously[1,2] in MACS buffer (PBS supplemented with 0.5% bovine serum albumin (BSA) and 2 mM EDTA). Cell subsets were then isolated by MACS following the manufacturer's protocols (Miltenyi Biotec). For example, to isolate $Gr1^+$ myeloid cells, scaffold cells were first incubated with FcR blocking reagents for 10 min, followed by incubating with anti-Gr1-biotin for another 10 min at 4 ºC. After washing with MACS buffer once, cell suspension was further incubated by anti-biotin microbeads and washed with MACS buffer again. Then the cell suspension was added to LS column (Miltenyi Biotec) in the magnetic field to separate the unlabeled $Gr1^-$ cells from magnetically labeled $Gr1^+$ myeloid cells. The kits or microbeads used in this study for isolating different subsets of lung or scaffold cells included CD45 microbeads, myeloid-derived suppressor cell isolation kit, anti-F4/80 microbeads ultrapure, CD11c microbeads ultrapure, CD90.2 microbeads, CD19 microbeads, CD49b microbeads, CD31 microbeads, and CD140a (PDGFRα) microbead kit.

**Condition media preparation.** To prepare scaffold or lung conditioned media from a tissue or a subset of cells enriched by MACS, single cell suspension were washed in PBS once and re-suspended in serum-free cell culture media (phenol red-free RPMI-1640 medium) at a concentration of $\sim5\times10^6$ cells/ml. Media were conditioned for 24 h at 37 ºC, after which the cells were removed by centrifugation. Total protein concentration of the conditioned media was quantified by a BCA protein assay (Thermo Fisher Scientific) and adjusted to 0.3 mg/ml for all *in vitro* studies.

***In vitro* extravasation of cancer cells.** $4\times10^4$ HUVEC cells in complete cell medium were seeded in the upper chamber of the Transwell system (6.5 mm diameter, 8 µm pore, Corning) and allowed to grow for two days to form a confluent monolayer. The HUVEC monolayer was activated by 20 ng/ml TNFα for 4 h and then rinsed with PBS three times. Cancer cells were stained with 20 µM CellTracker Green CMFDA dye in serum free medium at 37 ºC for 30 min and then recovered in complete cell culture medium for 1 h. Cancer cells were then trypsized, pelleted, and re-suspended in serum-free cell culture medium. $4\times10^4$ CMFDA-stained cancer cells in 100 µl of serum free medium were added to the upper chamber and 600 µl of conditioned medium, or 20 ng/ml S100A8/A9 chemoattractant (positive control), or serum-free cell culture medium (negative control) was added to the lower chamber. Two days later, the non-transmigrated cancer cells (TEM- cells) from the upper chamber were removed by a PBS soaked cotton swab. Transmigrated cancer cells (TEM+ cells) on the lower side of the filter were fixed and subsequently imaged by fluorescence microscopy (Zeiss) and quantified with ImageJ. In the antibody inhibition experiment, conditioned media were pre-incubated with 5 µg/ml anti-S100A8/A9 antibody at 37 ºC for 1 h prior to being loaded to the lower chamber. Each experiment was performed at least three times.

**Immunostaining and flow cytometry.** Lungs and scaffolds were retrieved at day 14 post orthotopic tumor inoculation and then processed to a single cell suspension as described previously.[1,2] Cells were first stained by a Live/Dead fixable blue stain (Thermo Fisher Scientific) for 15 min at 4 ºC, blocked by anti-mouse CD16/32 antibody (Biolegend), and then stained by antibody cocktails for 30 min at 4 ºC. When the immunostaining was finished, cells were washed with PBS, fixed by 4% paraformaldehyde for 15 min at room temperature and then re-suspended in ice-cold PBS for measurement by MoFlo Astrios #2 Cell Sorter. Four groups of antibody cocktails were separately used to stain the cells. The first group was designed to analyze leukocytes, myeloid cells, dendritic cells, monocytes, and macrophages, and this group included Pacific Blue anti-mouse Ly6-G/Ly6-C (Gr-1) antibody (Clone RB6-8C5, #108430, BioLegend), V500 rat anti-mouse CD11b antibody (Clone M1/70, #562128, BD Bioscience), PE/Cy7 anti-mouse F4/80 antibody (Clone BM8, #123114, BioLegend), APC anti-mouse CD11c antibody (Clone N418, #17-0114-82, eBioscience), and Alexa Fluor 700 (AF700) anti-mouse CD45 antibody (Clone 30-F11, #103128, BioLegend). The second group was designed to characterize T cells, B cells, and natural killer (NK) cells, and it included Pacific Blue anti-mouse CD19 antibody (Clone 6D5, #115526, BioLegend), V500 rat anti-mouse CD4 antibody (Clone RM4-5, #560783, BD Bioscience), FITC anti-mouse CD8a antibody (Clone 53-6.7, #100705, BioLegend), PE/Cy7 anti-mouse CD49b (pan-NK cells) antibody (Clone DX5, #108921, BioLegend), APC anti-mouse CD90.2 antibody (Clone 30-H12, #105311, BioLegend), and AF700 anti-mouse CD45 antibody. The third group was designed to distinguish endothelial cells and fibroblasts, and it included PE anti-mouse CD31 antibody (Clone 390, #102407, BioLegend), APC anti-mouse CD140a antibody (Clone APA5, #135907, BioLegend), and AF700 anti-mouse CD45 antibody. The fourth group was designed to characterize granulocytic myeloid cells and monocytic myeloid cells, and this group included Pacific Blue anti-mouse Gr1 antibody, V500 anti-mouse CD11b antibody, FITC anti-mouse Ly6-C antibody (Clone HK1.4, #128005, BioLegend), APC anti-mouse Ly-6G antibody (Clone 1A8, #127613, BioLegend), and AF700 anti-mouse CD45 antibody. All the antibodies were diluted to the manufacturer's recommended concentration for immunostaining.

**Quantitative RT-PCR (qRT-PCR).** Cell subsets from scaffolds and lungs were collected immediately after MACS cell separation for RNA extraction. Total RNA was extracted using PureLink RNA mini kit (Thermo Fisher Scientific) and cDNA was then synthesized using iScript$^{TM}$ cDNA synthesis kit (Bio-Rad). qRT-PCR was performed in 96-well plates on CFX Connect Real-Time System (Bio-Rad) with the following program: (1) 1 cycle of 95°C for 10 min; (2) 40 cycles of 95°C for 10 s and 58°C for 45 s. A total of 10 μl of reaction mixture was measured in each well, which included 5 μl of 2×SYBR Green supermix (Bio-Rad), 2 μl of primer mixture (2 μM of forward primers and 2 μM of reverse primers), 2 μl of cDNA (50 ng/ml) and 1 μl water. Three replicates were prepared for each sample. Data analysis was conducted using the ΔΔCt method in which all Ct values in each sample were first normalized to GAPDH. Each experiment was repeated at least three times.

**ELISA.** ELISA was used to identify the concentration of S100A8/A9 protein complex in the conditioned media. Briefly, 96-well ELISA plates were coated with 4 μg/ml of mouse S100A8/A9 heterodimer capture antibody overnight at room temperature. The plates were then blocked with 1% bovine serum albumin (BSA) in PBS for 1 h, followed by incubation with diluted conditioned media or standards for 2 h at room temperature. Subsequently, the plates were incubated with 40 ng/ml of mouse S100A8/A9 heterodimer detection antibody for another 2 h, followed by incubation with Streptavidin-HRP for 20 min and substrate solution for another 20 min. Copious washing by PBS was performed between incubations and all the incubations were performed at room temperature. The absorbance of each well at 650 nm with wavelength correction at 540 nm was recorded by a plate reader (BioTek Synergy H1). The chemoattractant concentration was calculated according to the standard curve and then normalized by the total protein concentration in the conditioned media. Three replicates were prepared for each sample and each experiment was repeated three times.

**Isolation of transmigrated cancer cells from Boyden chamber.** 4×10$^5$ HUVECs in complete medium were seeded in the upper chamber of a Transwell system (24.5 mm, 8μm pore, Corning) and grown for two days to be confluent on top of the porous filter. The HUVEC monolayer was activated by 20 ng/ml TNFα for 4 h, and then rinsed with PBS three times. 4×10$^5$ murine cancer cells in 1 ml serum-free medium were added to the upper chamber, and meanwhile, 2 ml of conditioned medium or 20 ng/ml S100A8/A9 or 100ng/ml CXCL12 or 40 ng/ml CCL2 in serum-free medium or medium supplemented with 0.1% or 1% FBS were added to the lower chamber. Two days later, the cells in the upper (non-transmigrated TEM- cells) and lower chamber (transmigrated TEM+ cells) were detached by cell dissociation buffer (enzyme-free, Gibco). The TEM- cells from the upper chamber were separated from HUVECs by MACS using a mouse cell depletion kit (Miltenyi Biotec). Briefly, cells collected from the upper chamber were incubated with mouse cell depletion cocktails for 15 min at 4 ºC, and then the magnetically labeled murine cancer cells were separated from unlabeled HUVECs using a LS column in the magnetic field. The ratio of TEM+ cancer cells vs. TEM- cancer cells was controlled to be less than 15% after two days' induction by changing the concentration of chemoattractants or conditioned media. This is because when the ratio of TEM+ cells to TEM- cells is high, the gene signature of TEM+ cancer cells can be diluted and covered by less aggressive cancer cells (Supplementary Fig. 18).

**Isolation of metastatic cancer cells from orthotopic cancer model.** Metastatic MDA-MB-231BR (231BR) human breast cancer cells in the lung and scaffold implants were isolated and expanded as described before.[3] Briefly, 2×10$^6$ 231BR cells in 50 μl PBS were inoculated into the fourth right mammary fat pad of 10-week-old female NSG mice at two weeks post scaffold implantation. At four weeks post tumor inoculation, mice were euthanized and then the primary tumor, scaffolds and lungs were retrieved and prepared to single cell suspension as described previously.[1,2] Human breast cancer cells were then isolated from murine tissue cells by MACS with a mouse cell depletion kit (Miltenyi Biotech) as described above. The human cell fraction were cultured in DMEM with 10% FBS and penicillin/streptomycin until growth of tumor cell colonies was evident. Following the first passage of cells, culture was continued without antibiotics. For all experiments the cell passage number was less than ten.

***in vitro* viability of 4T1/tdTomato/Luc cancer cells.** 4T1/tdTomato/Luc cancer cells were seeded to 96-well plate at a density of 800 cells/well in complete media supplemented with 0.6 mM D-luciferin, and then incubated at 37 ºC overnight to allow attachment. The next day, scaffolds and lungs that were retrieved from tumor-bearing BALB/c mice were prepared to single cell suspension as described before,[1,2] and Gr1$^+$ myeloid cell subset was isolated by MACS. These cells were subsequently added to 4T1/tdTomato/Luc tumor cell cultures in D-luciferin-included complete media and co-cultured for two days at 37 °C. The number of viable cancer cells was quantified by measuring the bioluminescence of 4T1/tdTomato/Luc cancer cells using a Synergy H1 hybrid multi-mode reader (BioTek). To measure the apoptotic cells, 4T1/tdTomato/Luc cells were incubated with scaffold or lung total cells in 96-well plates in complete cell culture media as described above. Two days later, suspended and weakly adherent cells were washed out by PBS and the adherent cancer cells were then trypsined and centrifuged at 400 rcf for 5 min at 4 °C and washed twice with PBS. Then 1×10$^5$ cancer cells were suspended in 100 μl of 1×binding buffer with 5 μl of AF488-conjugated Annexin V. Cells were incubated with Annexin V for 15 min at room temperature, and then centrifuged and wash once with PBS. Cells were fixed, resuspended in 200 μl of 1×binding buffer and measured by flow cytometer (Bio-Rad ZE5 Cell Analyzer).

***in vivo* bioluminescence imaging.** Female BALB/c mice (8 mice per group, 48 mice total) were subcutaneouly implanted with two 12 mm PCL scaffolds on the upper back (implant groups, n = 24) or received mock surgery (control groups, n =24). 14 days later, 2×10$^6$ 4T1/tdTomato/Luc cancer cells in 50 μl of PBS were inoculated into the fourth right mammary fat pad by injection. At days 10, 15 and 20 after tumor inoculation, primary tumor were resected as described previously.[1] Immediately after surgery, the mice were intraperitoneally injected with 150 μl of 63 mM D-luciferin solution and bioluminescence imaging was conducted with an IVIS (PerkinElmer) to observe lung metastasis. The absence of the resected primary tumor allowed for detection of lung metastases that would otherwise be obscured by the very bright primary tumor signal.

**Data analysis.** Data are presented as standard error of the mean (SEM). Animal studies were performed with at least two independent replicates of three to eight female BALB/c mice per group. *p*-values were determined using Student's *t* test for single comparison when not otherwise specified. We note that our bioluminescence data was not normally distributed (Fig. 5D), so we used the non-parametric Mann Whitney test to assess *p*-values. We used the Fisher's exact test to analyze our metastasis *vs.* no metastasis categorical data (Fig. 5D). Here we note that different mice were resected on days 10, 15, and 20 post tumor inoculation (48 mice total in experiment, 24 mock surgery mice, 24 scaffold implanted mice), so we can treat each mouse on each day as independent for purposes of pooling and statistical comparison.

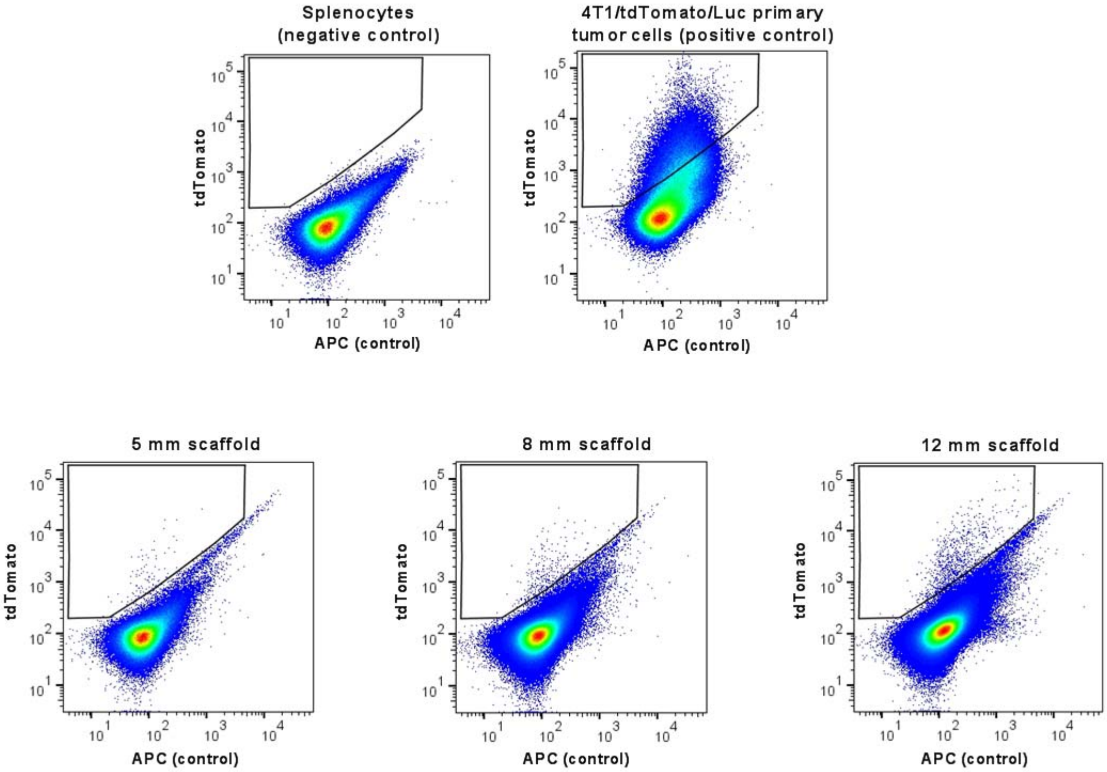


**Supplementary Fig. 1. Representative density plots for tdTomato-positive 4T1 cancer cells that metastasized from the primary tumor to the scaffold implants with different sizes in orthotopic injection spontaneous metastasis model.** Splenocytes and 4T1/tdTomato/Luc primary tumor cells were used as negative control and positive control, respectively, in the flow cytometric analysis. Female BALB/c mice were orthotopically inoculated with $2\times10^6$ of 4T1/tdTomato/Luc cancer cells 14 days after scaffold implantation. 14 days after tumor inoculation, mice were euthanized and single cell suspensions were prepared from scaffolds and tissues. Single cells from one scaffold were diluted in 500 µl PBS and then 100 µl was measured by flow cytometer.

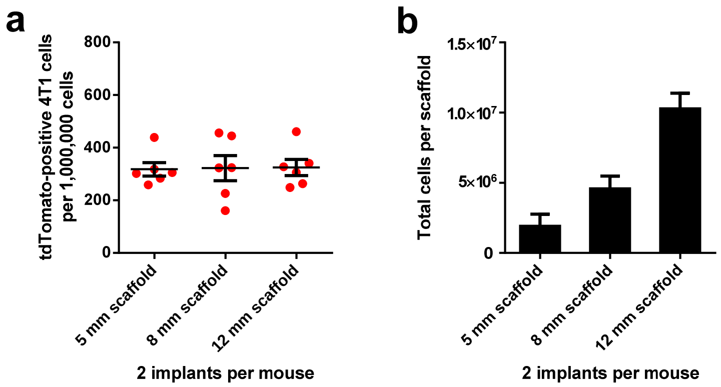


**Supplementary Fig. 2. Relative abundance of tdTomato-positive 4T1 cancer cells to total cells in scaffold implants with different sizes (a) and total cells per scaffold (b).** PCL scaffolds (5 mm, 8 mm or 12 mm in diameter) were subcutaneously implanted to female BALB/c mice (2 implants per mouse), and two weeks later, $2\times10^6$ of 4T1/tdTomato/Luc cells were orthotopically inoculated to the forth mammary fat pad of mice. At day 14 after tumor inoculation, scaffolds were retrieved and prepared to single cell suspension, and the fluorescent tdTomato-positive 4T1 cancer cells that spontaneously metastasized from primary tumor to scaffolds were quantified by flow cytometer.

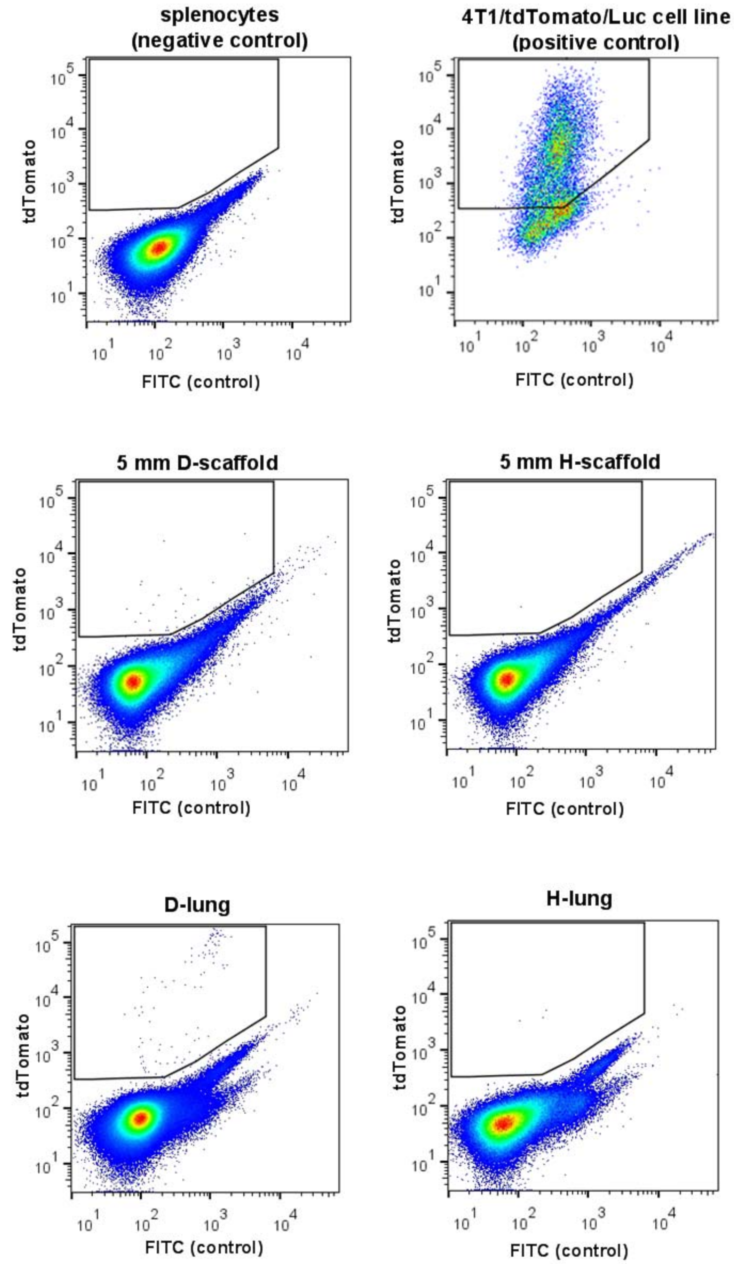


**Supplementary Fig. 3. Representative density plots for tdTomato-positive 4T1 cancer cells that extravasated to 5 mm scaffold implants and lungs in intracardiac injection metastasis model.** Splenocytes and 4T1/tdTomato/Luc cell line were used as negative control and positive control, respectively, in the flow cytometric analysis. Female BALB/c mice were orthotopically inoculated with $2\times10^6$ of unlabeled 4T1 cancer cells or injected with PBS (control) at two weeks post scaffold implantation. At day 14 post tumor inoculation, $1\times10^6$ of fluorescent 4T1/tdTomato/Luc cells were intracardiacally injected to tumor-bearing mice ("diseased" D) and tumor-free mice ("healthy" H). And one day later, tissues were retrieved and prepared to single cell suspension, and tdTomato-positive cancer cells in scaffold implants and lungs were measured by flow cytometer.

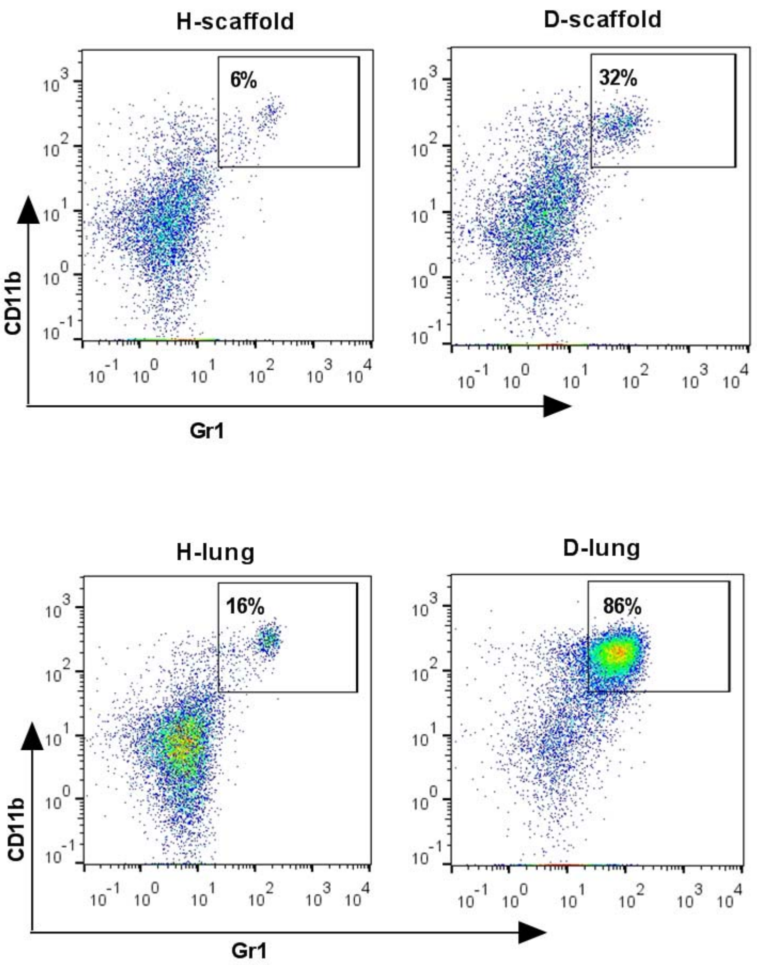


**Supplementary Fig. 4. Representative density plots for CD11b$^+$Gr1$^+$ myeloid cells in scaffold implants and lungs.** Scaffold implants and lungs were retrieved from mice bearing 4T1 tumor ("diseased" D) or tumor-free mice ("healthy" H) at two weeks post tumor inoculation.

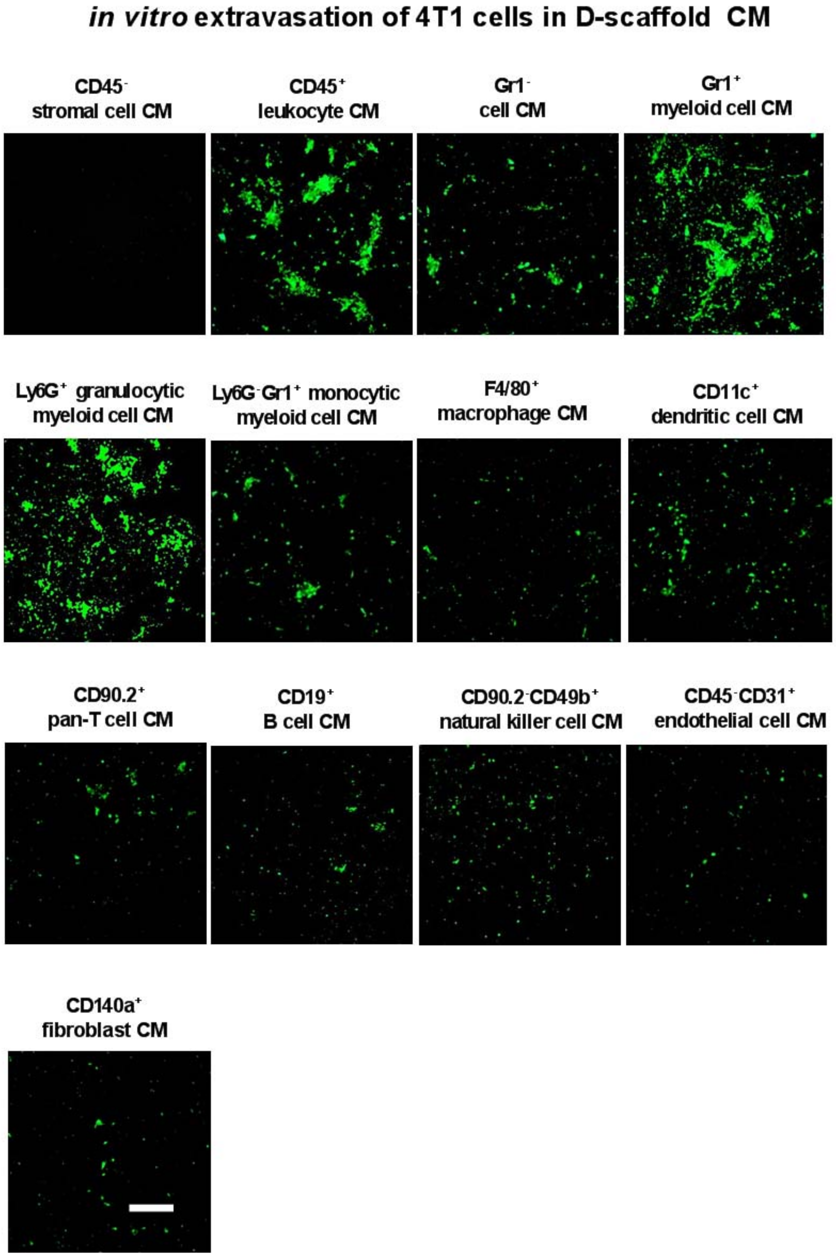


**Supplementary Fig. 5. Microscopy images of 4T1 cancer cells that transmigrated towards conditioned media prepared from different diseased scaffold cell subsets**. The surface markers listed above each image indicate the antibodies used in magnetic-activated cell sorting. 4T1 cancer cells were allowed to extravasate through a HUVEC monolayer and transmigrate through a Boyden chamber for two days before imaging. Scale bar = 150 µm. D-scaffold: diseased scaffold retrieved from tumor-bearing mice; CM: conditioned media.

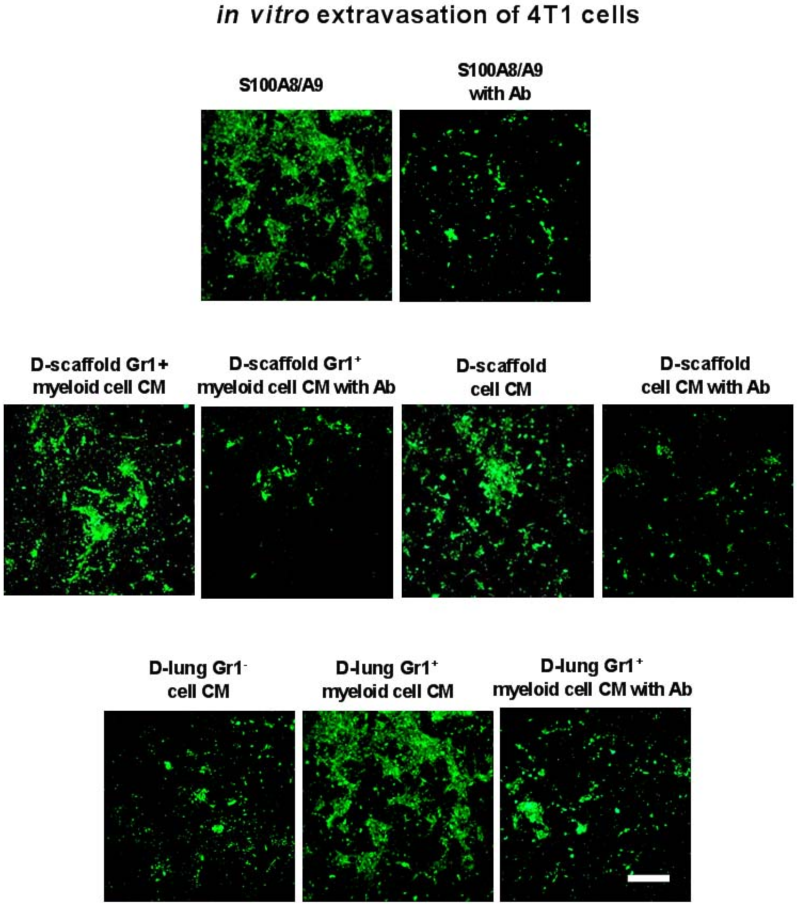


**Supplementary Fig. 6. Microscopy images of 4T1 cancer cells that transmigrated towards 20 ng/ml S100A8/A9 (positive control) or conditioned media in the absence or presence of anti-S100A8/A9 antibody.** D-scaffold and D-lung indicate that tissues were retrieved from tumor-bearing mice (diseased). 4T1 cancer cells were allowed to extravasate through a HUVEC monolayer and transmigrate through a Boyden chamber for two days before imaging. Scale bar = 150 μm. CM: conditioned media; Ab: anti-S100A8/A9 antibody.

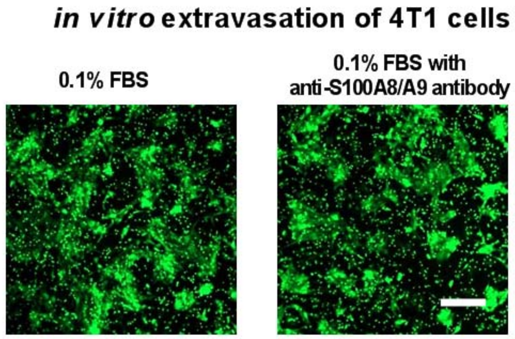


**Supplementary Fig. 7. Anti-S100A8/A9 antibody does not inhibit FBS-induced 4T1 cell transmigration.** 4T1 cancer cells were allowed to extravasate through a HUVEC monolayer and transmigrate through a Boyden chamber towards 0.1% FBS for two days before imaging. Scale bar = 150 µm. FBS: fetal bovine serum.

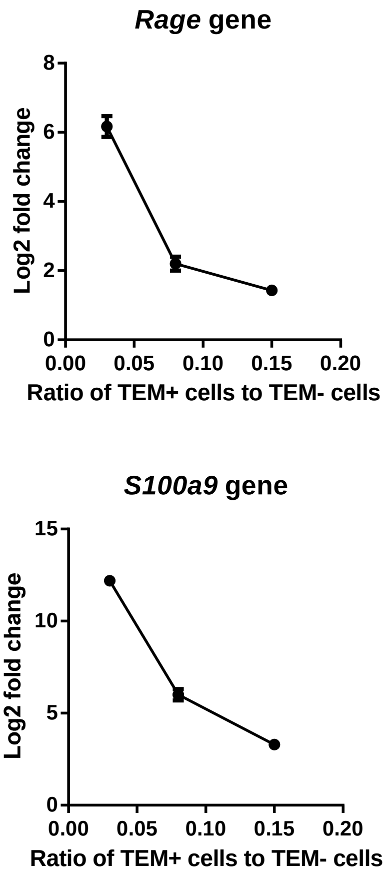


**Supplementary Fig. 8. The initiating 4T1 cancer cells that transmigrated toward D-lung conditioned media overexpressed *Rage* and *S100a9* genes relative to later transmigrating cells**. Transmigration of 4T1 cancer cells was induced by D-lung conditioned media in an *in vitro* extravasation model for 6 h, 24 h and 48 h, respectively, resulting in the ratio of TEM+ cell number to TEM- cell number to be ~3%, ~8%, and ~15%, respectively. The relative expression of *Rage* and *S100a9* genes between TEM+ cells and TEM- cells was compared at different time points. TEM: transendothelial migration. D-lung: diseased lung retrieved from tumor-bearing mice.

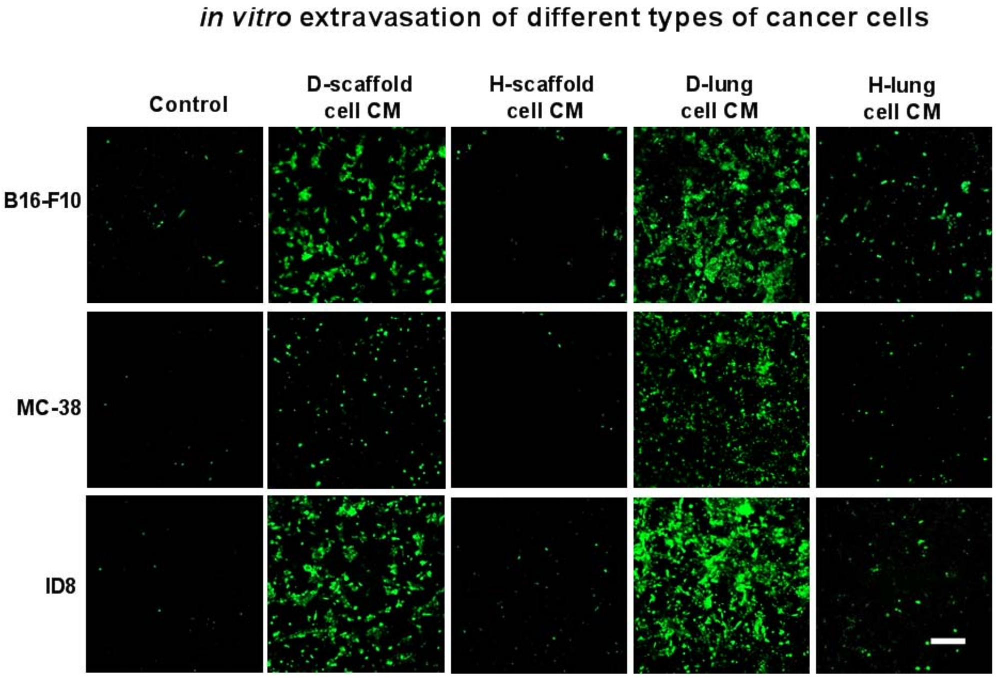


**Supplementary Fig. 9. Microscopy images of transmigrated B16-F10 murine melanoma cells, MC-38 murine colon cancer cells and ID8 murine epithelial ovarian cancer cells**. Transmigration was induced by conditioned media prepared from scaffolds and lungs retrieved from tumor-bearing mice (“diseased” D) or tumor-free mice (“healthy” H). Cancer cells were allowed to extravasate through a HUVEC monolayer and transmigrate through a Boyden chamber towards conditioned media for two days before imaging. Scale bar = 150 µm. CM: conditioned media.

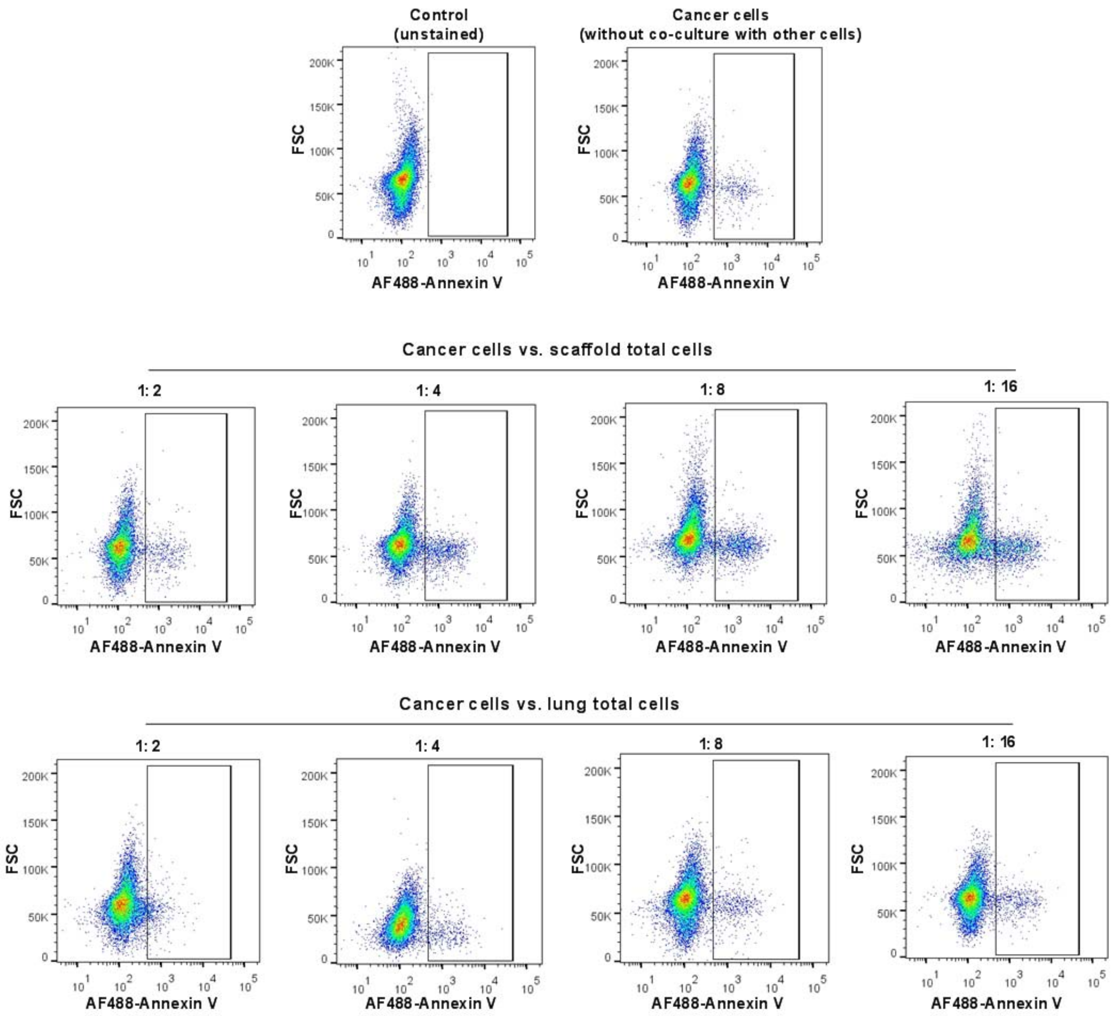


**Supplementary Fig. 10. Representative density plots for apoptotic cell subpopulation in 4T1 cancer cells co-cultured with scaffold or lung total cells.** 4T1/tdTomato/Luc cancer cells were co-cultured with total cells derived from D-scaffold or D-lung at different ratios for two days in complete cell culture media. And then scaffold or lung cells were washed out by PBS, and 4T1 cells were trypsined, stained by AF488-labeled Annexin V, fixed and then measured by flow cytometry. D: diseased.

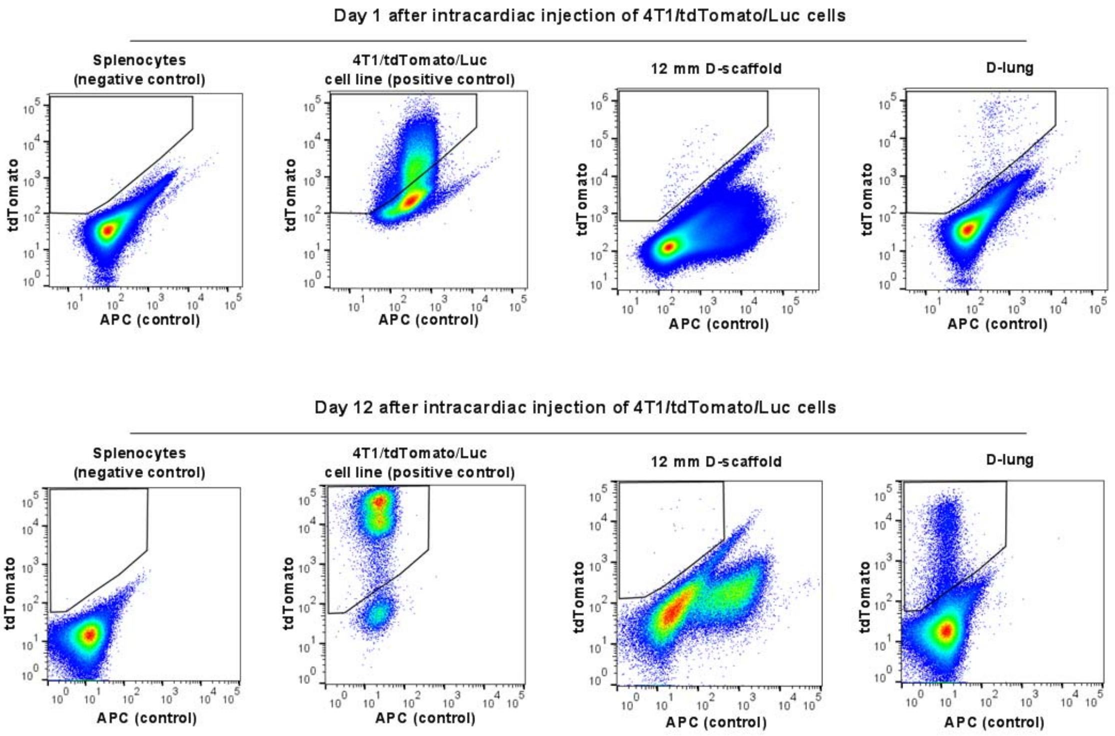


**Supplementary Fig. 11. Representative density plots for tdTomato-positive 4T1 cancer cells that extravasated to 12 mm scaffold implants and lungs at 1 and 12 days after intracardiac injection of 4T1/tdTomato/Luc cells in mice bearing unlabeled 4T1 tumor.** Scaffold-bearing female BALB/c mice were first inoculated with $2\times10^6$ unlabeled 4T1 cells by orthotopic injection, and 14 days later, the tumor-bearing mice received $1\times10^6$ 4T1/tdTomato/Luc cells by intracardiac injection. The 4T1 cells introduced by intracardiac injection were cleared from circulation in one or two days[6]. tdTomato-labeled 4T1 cells that extravasated to the scaffold failed to proliferate and died as they are less abundant 12 days after intracardiac injection than 1 day. In contrast, tdTomato-labeled 4T1 cells that extravasated from circulation to the lung proliferated and are much more abundant on 12 days after intracardiac injection than 1 day.

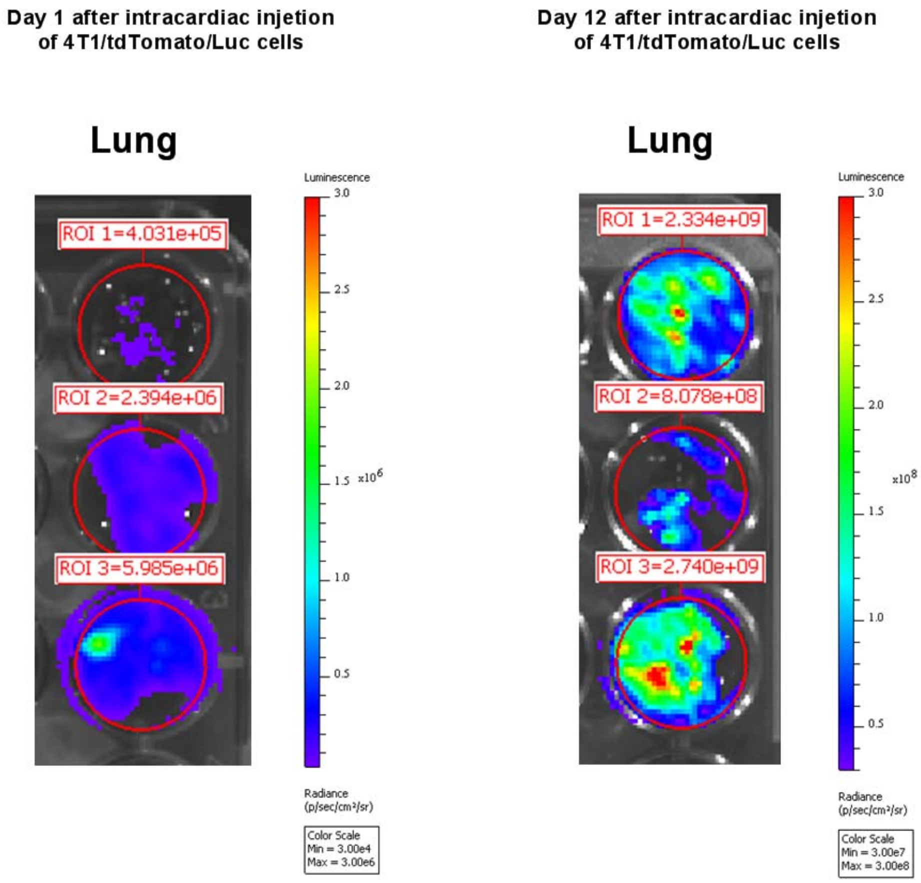


**Supplementary Fig. 12. Bioluminescence imaging of lungs at 1 day and 12 days after intracardiac injection of 4T1/tdTomato/Luc cells in mice bearing an unlabeled 4T1 tumor**. Female BALB/c mice were orthotopically inoculated with $2\times10^6$ unlabeled 4T1 cancer cells two weeks after scaffold implantation (two 12 mm scaffolds per mouse). 14 days later, $1\times10^6$ 4T1/tdTomato/Luc cells were introduced into circulation of the tumor-bearing mice by intracardiac injection. 1 and 12 days after intracardiac injection, lungs were retrieved and incubated in 1.2 mM D-luciferin solution for 10 min on ice, and then imaged. Total flux for the lungs of 3 mice is shown for each condition.

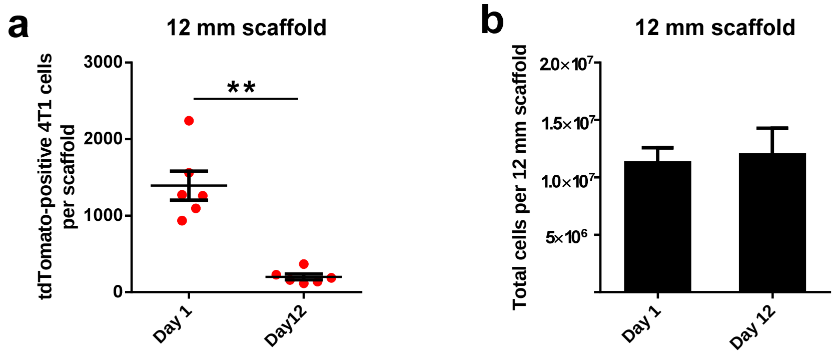


**Supplementary Fig. 13. tdTomato-positive 4T1 cells per scaffold (a) and total cells per scaffold (b) at 1 and 12 days after intracardiac injection of 4T1/tdTomato/Luc cells in mice bearing unlabeled 4T1 tumor.** Female BALB/c mice were orthotopically inoculated with $2\times10^6$ unlabeled 4T1 cancer cells at two weeks after scaffold implantation (two 12 mm scaffolds per mouse). 14 days later, $1\times10^6$ 4T1/tdTomato/Luc cells were introduced into circulation of tumor-bearing mice by intracardiac injection. And 1 and 12 days after intracardiac injection, tissues were retrieved and processed into single cell suspensions. tdTomato-positive cancer cells in scaffold implants and lungs were measured by flow cytometer. $^{**}p < 0.01$, from student *t*-test.

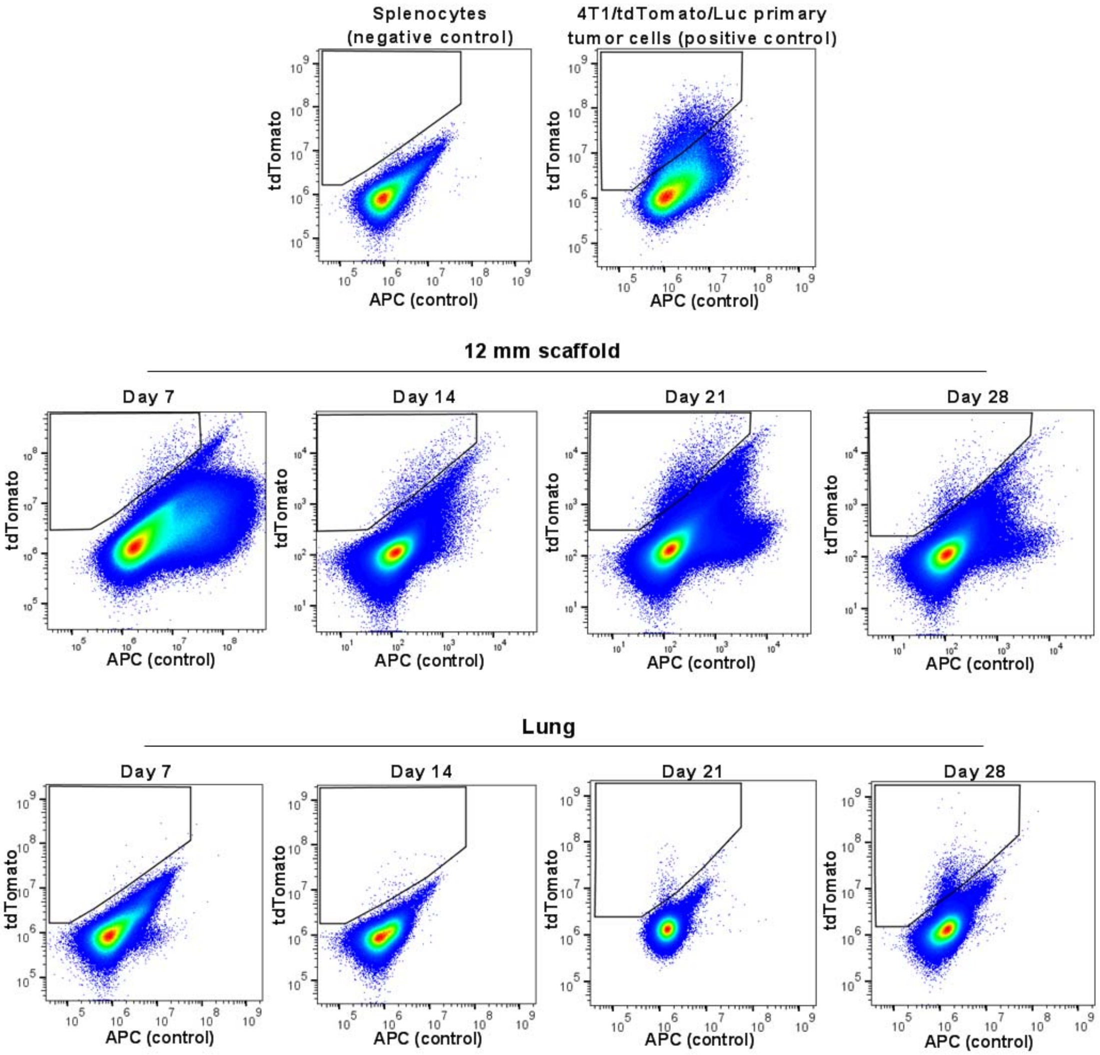


**Supplementary Fig. 14. Representative density plots for tdTomato-positive 4T1 cancer cells that metastasized to scaffold implants and lungs at different time points post orthotopic inoculation of 4T1/tdTomato/Luc cells.** Splenocytes and 4T1/tdTomato/Luc primary tumor cells were used as negative control and positive control, respectively, in the flow cytometric analysis. Female BALB/c mice were orthotopically inoculated with $2\times10^6$ of 4T1/tdTomato/Luc cancer cells at two weeks post scaffold implantation (two 12 mm scaffolds per mouse). At days 7, 14, 21 and 28 post tumor inoculation, tissues were retrieved and prepared to single cell suspension, and tdTomato-positive cancer cells in scaffold implants and lungs were measured by flow cytometer.

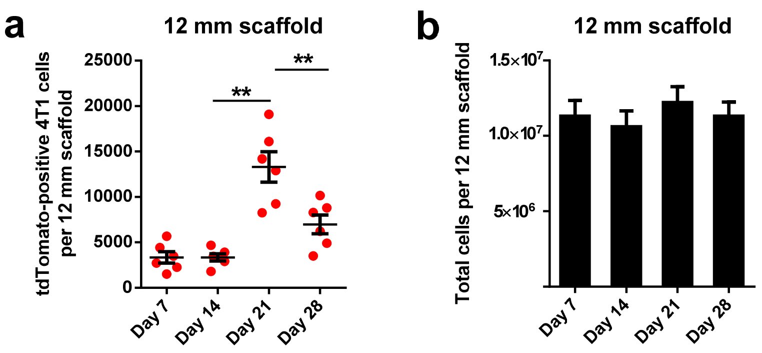


**Supplementary Fig. 15. tdTomato-positive 4T1 cells per scaffold (a) and total cells per scaffold (b) at days 7, 14, 21 and 20 after orthotopic inoculation of 4T1/tdTomato/Luc cells.** Female BALB/c mice were orthotopically inoculated with $2\times10^6$ of 4T1/tdTomato/Luc cancer cells at two weeks post scaffold implantation (two 12 mm scaffolds per mouse). At days 7, 14, 21 and 28 post tumor inoculation, tissues were retrieved and prepared to single cell suspension, and tdTomato-positive cancer cells in scaffold implants and lungs were measured by flow cytometer. $^{**}p < 0.01$, from student *t*-test.

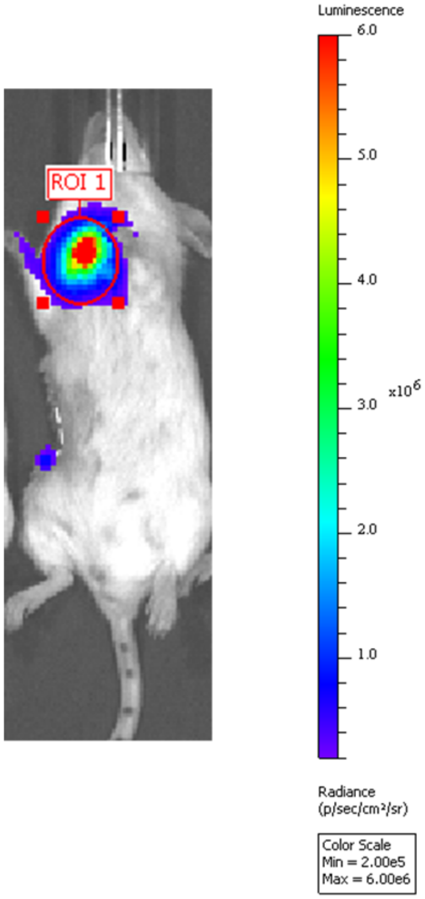


**Supplementary Fig. 16. The ROI area used to calculate the total flux of lung metastasis in Fig. 5D.** The ROI with a fixed size was placed to the brightest area in lungs for each mouse.

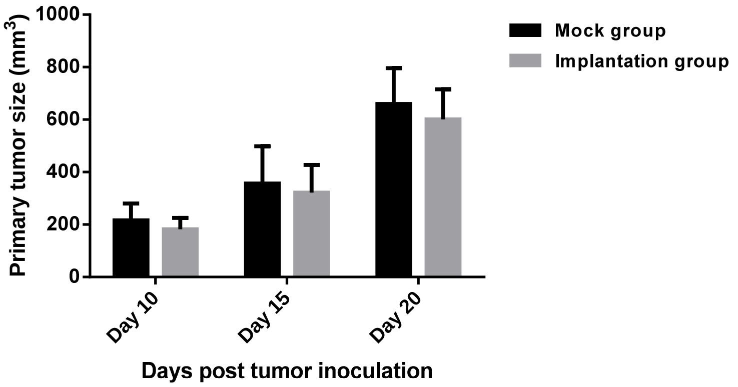


**Supplementary Fig. 17. Averaged primary tumor size at days 10, 15 and 20 after tumor inoculation.** The implantation group of mice (n = 24) was implanted with 12 mm PCL scaffolds (2 implants per mouse), while the mock group of mice received mock surgery (control, n = 24). 14 days after implantation/mock surgery, both groups of mice were orthotopically inoculated with $2\times10^6$ of 4T1/tdTomato/Luc cancer cells.

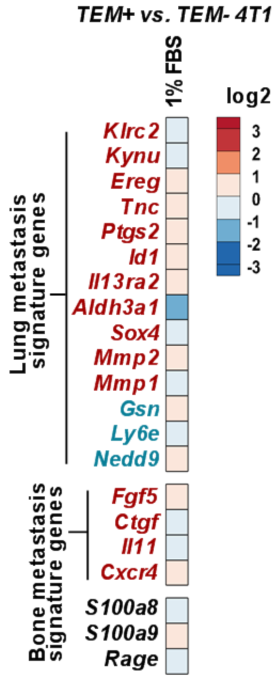


**Supplementary Fig. 18. A high ratio of TEM+ cells to TEM- cells obscured their gene expression difference.** Heatmap of expression of organotropism signature genes in 1% FBS-induced transmigrated 4T1 (TEM+ cells) *vs*. non-transmigrated 4T1 (TEM- cells) isolated from Boyden chamber. In the heatmap, red genes and blue genes represent overexpressed and underexpressed, respectively, organotropism signature genes in metastatic cancer cells relative to parental cells reported in the literature.[4,5] Transmigration of cancer cells was induced for two days, and the ratio of TEM+ cell number to TEM- cell number was > 50% in this condition. TEM: transendothelial migration. FBS: fetal bovine serum

**References for SI reference citations**


1. Rao, S. S. *et al*. Enhanced survival with implantable scaffolds that capture metastatic breast cancer cells *in vivo*. *Cancer Res* **76**, 5209-5218 (2016).
2. Azarin, S. M. *et al*. *In vivo* capture and label-free detection of early metastatic cells. *Nat Commun* **6**, 8094 (2015).
3. Bushnell, G. G. *et al*. Biomaterial scaffolds recruit an aggressive population of metastatic tumor cells *in vivo*. *Cancer Res* **79**, 2042-2053. (2019).
4. Minn, A. J. *et al*. Genes that mediate breast cancer metastasis to lung. *Nature* **436**, 518-524 (2005).
5. Kang, Y. *et al.* A multigenic program mediating breast cancer metastasis to bone. *Cancer Cell* **3**, 537-549 (2003).
6. Basse, P., Hokland, P., Heron, I. & Hokland, M. Fate of tumor cells injected into left ventricle of heart in BALB/c mice: role of natural killer cells. *J Natl Cancer Inst* **80**, 657-665 (1998).
